# Discovery of benzo[c]phenanthridine derivatives with potent activity against multidrug resistant *Mycobacterium tuberculosis*

**DOI:** 10.1101/2022.11.07.515485

**Authors:** Zhiqi Sun, Yi Chu Liang, Chen Lu, Andréanne Lupien, Zhongliang Xu, Stefania Berton, Marcel A. Behr, Weibo Yang, Jim Sun

## Abstract

*Mycobacterium tuberculosis* (Mtb), the pathogen responsible for tuberculosis (TB), is the leading cause of bacterial disease-related death worldwide. Current antibiotic regimens for the treatment of TB remain dated and suffer from long treatment times as well as the development of drug-resistance. As such, the search for novel chemical modalities that have selective or potent anti-Mtb properties remains an urgent priority, particularly against multidrug resistant (MDR) Mtb strains. Herein, we design and synthesize 35 novel <u>b</u>enzo[c]<u>p</u>henanthridine <u>d</u>erivatives (BPD). The two most potent compounds, BPD-6 and BPD-9, accumulated within the bacterial cell and exhibited strong inhibitory activity (MIC_90_ ∼ 2-10 μM) against multiple *Mycobacterium* strains, while remaining inactive against a range of other Gram-negative and Gram-positive bacteria. BPD-6 and BPD-9 were also effective in reducing Mtb viability within infected macrophages. The two BPD compounds displayed comparable efficacy to rifampicin, a critical frontline antibiotic used for the prevention and treatment of TB. Importantly, BPD-6 and BPD-9 inhibited the growth of multiple MDR Mtb clinical isolates, suggesting a completely novel mechanism of action compared to existing frontline TB dugs. The discovery of BPDs provides novel chemical scaffolds for anti-TB drug discovery.

**TOC/GRAPHICAL ABSTRACT:** 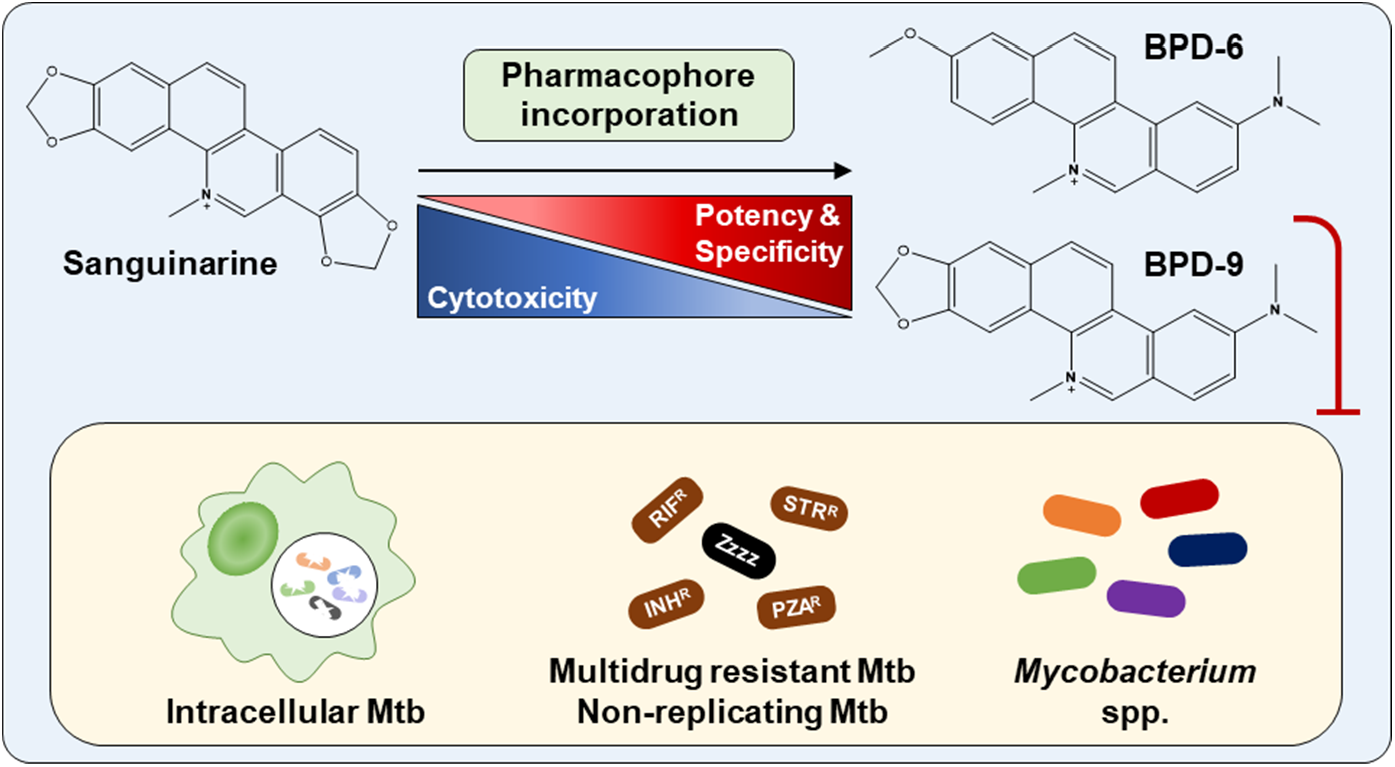

## INTRODUCTION

Tuberculosis (TB) has existed throughout human history and is still threatening millions of lives each year.^1^ Since the discovery of *Mycobacterium tuberculosis* (Mtb) in the 19^th^ century, the search for novel, specific and effective antibiotics against Mtb has remained a major priority for TB drug discovery. Unlike Gram-positive or Gram-negative bacteria, the waxy, mycolic acid rich cell wall of Mtb is the first barrier to overcome in TB drug discovery. Small chemical molecules that are not permeable through the Mtb cell wall need to be coated or encapsulated to increase penetration *in vivo* or *ex vivo*^2,3^, necessitating extra research efforts. Additionally, under the pressure of host immunity and antibiotics, Mtb can adapt via a non-replicating state that facilitates non-heritable resistance or tolerance to most conventional antibiotics.^4^ As such, efficacy of TB drug candidates should be assessed using *in vitro* and *in vivo* models of non-replicating Mtb (NR-Mtb). A number of *in vitro* models have been established to generate low-metabolism Mtb cultures, including hypoxia, nutrient starvation, low pH, and the combination of multiple stresses.^5^ Of the frontline TB drugs, rifampicin (RIF) is the most well characterized to inhibit NR-Mtb under acidic conditions.^6,7^ Another frontline TB drug, pyrazinamide, eliminates NR-Mtb *in vivo* and *in vitro* by targeting non-specific pathways.^8^ Furthermore, recently approved TB drugs, bedaquiline^9^ and pretomanid^10^ have demonstrated efficacy against NR-Mtb *in vivo*^6^.

To find new mechanisms of action and new chemical scaffolds, natural products possess a range of benefits for drug discovery.^3,11^ Natural products are ideal candidates for drug discovery due to their unique structural and scaffold diversity. Typically, natural products tend to be compounds with higher hydrophilicity and greater molecular rigidity than synthetic chemicals, which are favourable traits for drug development. Importantly, as most natural products are bioactive molecules refined by evolution to serve specific biological functions such as endogenous regulation and inter-organism competition, they are highly relevant to anti-cancer and anti-infection applications.^12^ Rifampicin is a successful example of a semisynthetic drug derived from the natural compound rifamycin.^13^ Other well-known examples include streptomycin, the first successful antibiotic against TB that was isolated from *Streptomyces griseus*, as well as the second-line TB antibiotics kanamycin, produced by *Streptomyces kanamycetius*, and its semi-synthetic analogue amikacin.^14^

Sanguinarine (SG), a natural benzo[c]phenanthridine alkaloid extracted from *Sanguinaria canadensis*, has been used as an antiseptic herbal essence and was used as an antimicrobial agent in toothpaste and mouthwash.^15,16^ SG has shown antibacterial effects against various Gram-positive and Gram-negative bacteria.^17–19^ In a recent study, SG has been shown to be an inhibitor of the 2-ketogluconate pathway, which is important for glucose metabolism in *Pseudomonas aeruginosa*.^19^ There have also been reports demonstrating the synergistic effects of SG and antibiotics.^17,20^ However, the antibacterial effects of SG against mycobacteria have not been explored and may represent a missed opportunity.

Herein, we designed and synthesized 35 unique derivatives and analogues of SG. Phenotypic activity screening against actively replicating Mtb identified 5 hits within this group of new compounds that possessed significantly improved anti-Mtb activity compared to SG, which surprisingly only showed modest activity against Mtb. The effectiveness of the two most potent compounds, BPD-6 and BPD-9, was characterized in detail in a series of experiments. We demonstrate that BPD-6 and BPD-9 possess low-micromolar inhibitory activity against multiple mycobacterial species, but were inactive against other Gram-negative and Gram-positive bacteria, showing unique specificity against mycobacteria. Importantly, both compounds exhibit reduced cytotoxicity, a known undesirable property of SG, and showed efficacy against Mtb within infected human macrophages. Importantly, BPD-6 and -9 was effective in inhibiting NR-Mtb. In addition, both compounds exhibited potent activity against multiple multidrug resistant (MDR) Mtb clinical isolates, which demonstrates a unique mechanism of action compared to existing frontline TB antibiotics. Collectively, our data demonstrate that novel benzo[c]phenanthridine derivatives have the potential to be developed into selective anti-mycobacterial drugs to provide new chemical options for TB antibiotic discovery.

## RESULTS AND DISCUSSION

### Generation of novel benzo[c]phenanthridine derivatives with antibacterial activity

Given that the antibacterial activity of SG has not been reported for *M. tuberculosis*, we first assessed whether SG possessed any inhibitory activity against Mtb mc^2^6206, an auxotrophic strain that retains a similar drug susceptibility profile as its virulent parent strain H37Rv.^21^ Using the resazurin microtiter growth inhibition assay (REMA)^22^, SG showed only modest activity against Mtb mc^2^6206, reaching a minimum inhibition concentration (MIC_90_) of 85 μM (Figure 1A). To define and improve the activity of the pharmacophore responsible for the anti-Mtb activity of SG, we used a pharmacophore incorporation strategy to design new benzo[c]phenanthridine derivatives. Three points of pharmacomodulation were applied to modify the structure of SG: 1) derivatives modified at rings A and D of the benzo[c]phenanthridine moiety; 2) simplification of the quaternary pyridinium skeleton of the phenanthridinium core; 3) dearomatizing ring C and introducing diversified functional groups at C-8 or the N atom (Figure 1B).

**Figure 1.**
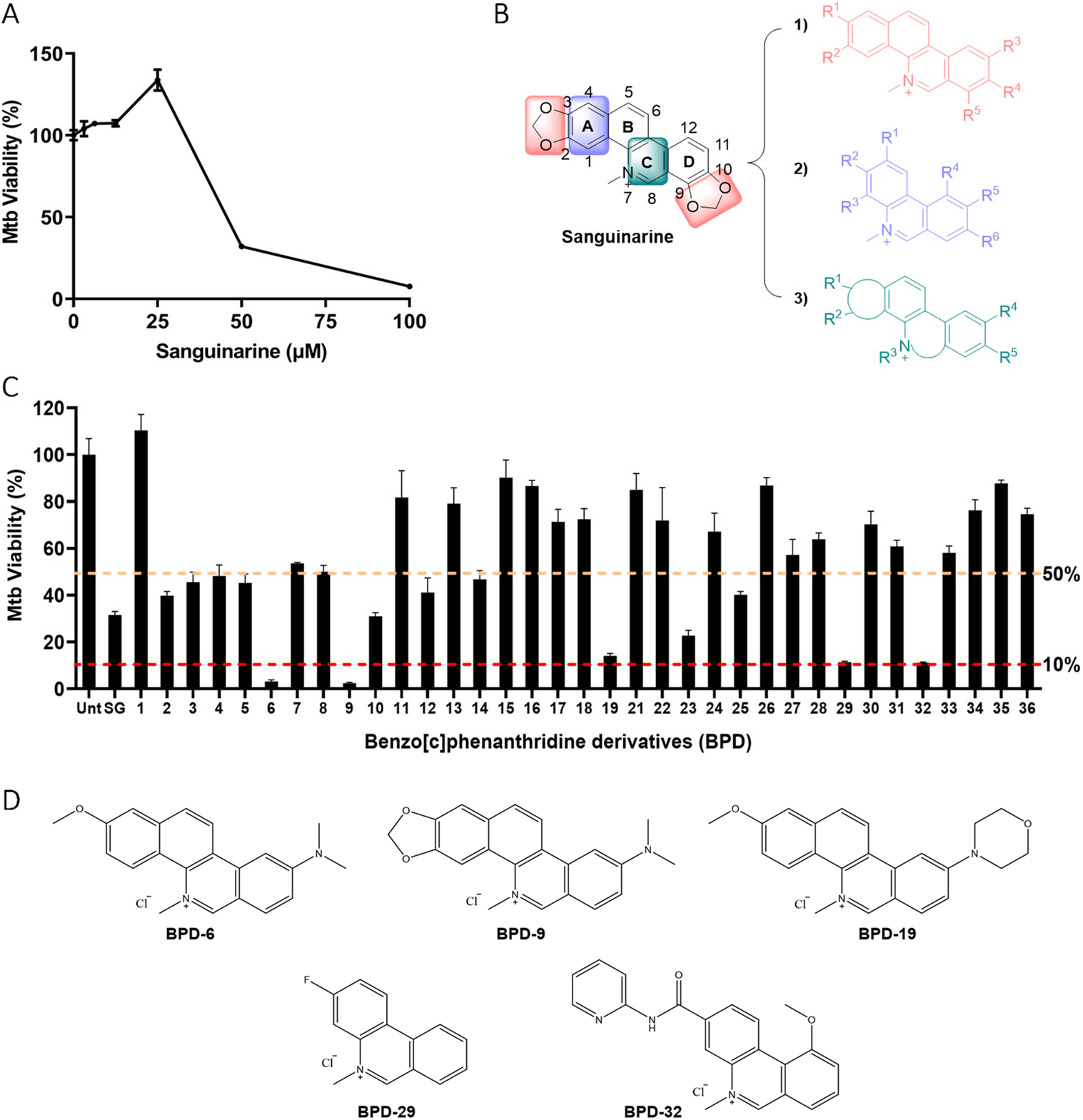
Generation of novel benzo[c]phenanthridine derivatives with antibacterial activity. (A) Activity of Sanguinarine (SG) against Mtb mc^2^6206 was determined using the REMA assay. Mtb viability is normalized to maximal bacterial growth in the absence of compounds as 100%. (B) Pharmacomodulation strategy to generate novel benzo[c]phenanthridine derivatives (BPD) from SG. (C) BPD compounds were screened for activity against Mtb mc^2^6206 at 40 μM using the REMA assay. Mtb viability is normalized to maximal bacterial growth in the absence of compounds as 100%. SG, sanguinarine; Unt, untreated. (D) Chemical structures of compounds with highest Mtb inhibitory activity. Data in (A, C) represent the mean ± SEM of three independent experiments.

In total, 35 new benzo[c]phenanthridine derivatives were synthesized and screened for anti-Mtb activity using the REMA assay. In the initial screen using a fixed concentration of 40 μM, 15 (43%) of the new compounds were able to inhibit Mtb growth by at least 50% compared to untreated controls (Figure 1C). Of these 15 hits, five compounds showed significantly improved activity compared to SG and inhibited the growth of Mtb by ≥ 85% (Figure 1C, D). The two most potent hits, compounds #6 and #9, hereafter designated as BPD-6 and BPD-9 (benzo[c]phenanthridine derivatives) (Scheme 1 and Figures S1, S2) showed greater than 90% inhibition and were thus selected for further characterization. Based on the structures of the active hits, the minimal pharmacophore of BPD derivatives are rings B-C-D. The azanium of ring C is critical for anti-Mtb activity, and substitutions on rings B and D affect the activity of the compounds. The benzo-segment of ring B could be simplified to generate derivatives with considerable activity, and additional hydrogen receptors or donors could be combined for a productive gain in activity, such as in BPD-29 and BPD-32. The dioxolane on ring D is dispensable for activity, and introducing any substituents on C-9 or C-10 on ring D resulted in a loss of activity, which may be due to a blocking interaction with the target. However, introduction of nitrogen-containing groups at C-11 of ring D improved the potency significantly (BPD-6, 9, and 19).

**Scheme 1.**
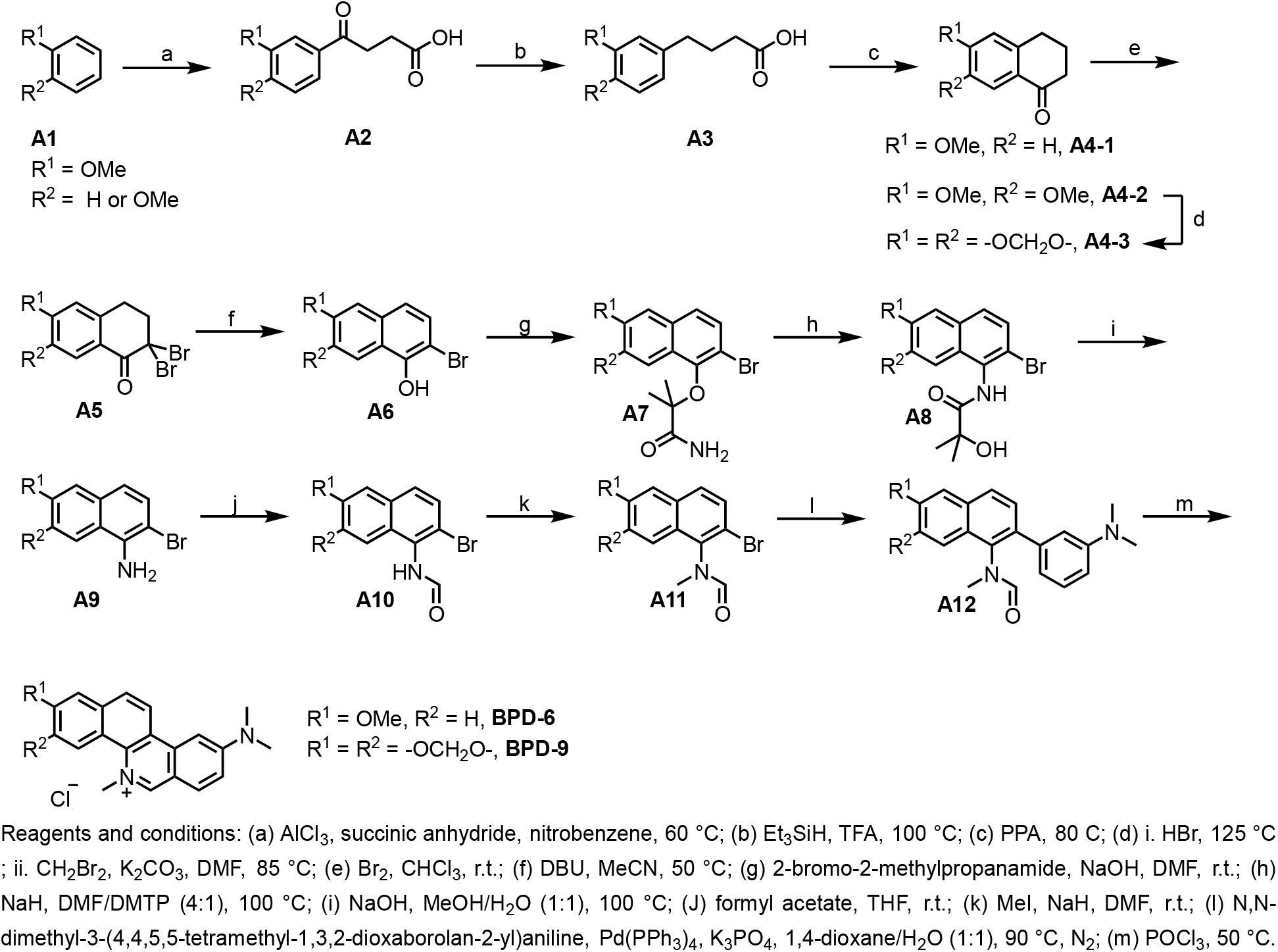
Synthetic route for BPD-6 and BPD-9.

### BPD-6 and BPD-9 significantly inhibit the growth of mycobacteria

The REMA growth inhibition assay was used to determine the dose-dependent inhibition activity of BPD-6, -9, -29, -32 against Mtb mc^2^6206 (Figure 2A). This assay showed that BPD-6 and BPD-9 possessed the most potent anti-Mtb activity, reaching an MIC_90_ of 10 μM and 6 μM, respectively (Table 1). The activity of BPD-6 and BPD-9 thus show an 8-fold and 14-fold improvement over SG, respectively, confirming that the new BPD derivatives possess significantly increased anti-Mtb activity. While SG is broadly active against several bacterial species, we wondered if this property was retained in our new compounds. To assess the specificity of BPD-6 and -9, we performed REMA growth inhibition assays on various mycobacterial strains. To our surprise, BPD-6 and -9 retained potent inhibitory effect against *Mycobacterium bovis* BCG, *Mycobacterium kansasii*, and *Mycobacterium smegmatis*, while showing no effect against other Gram-positive and Gram-negative bacteria (Table 1). The specificity of an antibiotic to a particular pathogenic genus or species may have major advantages for treatment potential since it is less likely to adversely affect the host microbiome.^23–25^ The specificity of BPD-6 and -9 suggest a possible mechanism or target of inhibition that exists specifically in mycobacteria. It has been reported that SG binds to FstZ in various bacterial species, including *E. coli*^26^ to inhibit their growth by preventing cell division.^18^ Given that FstZ is highly conserved across bacteria species, our results indicate that BPD-6 and -9 inhibit a unique target specific to mycobacteria.

**Figure 2.**
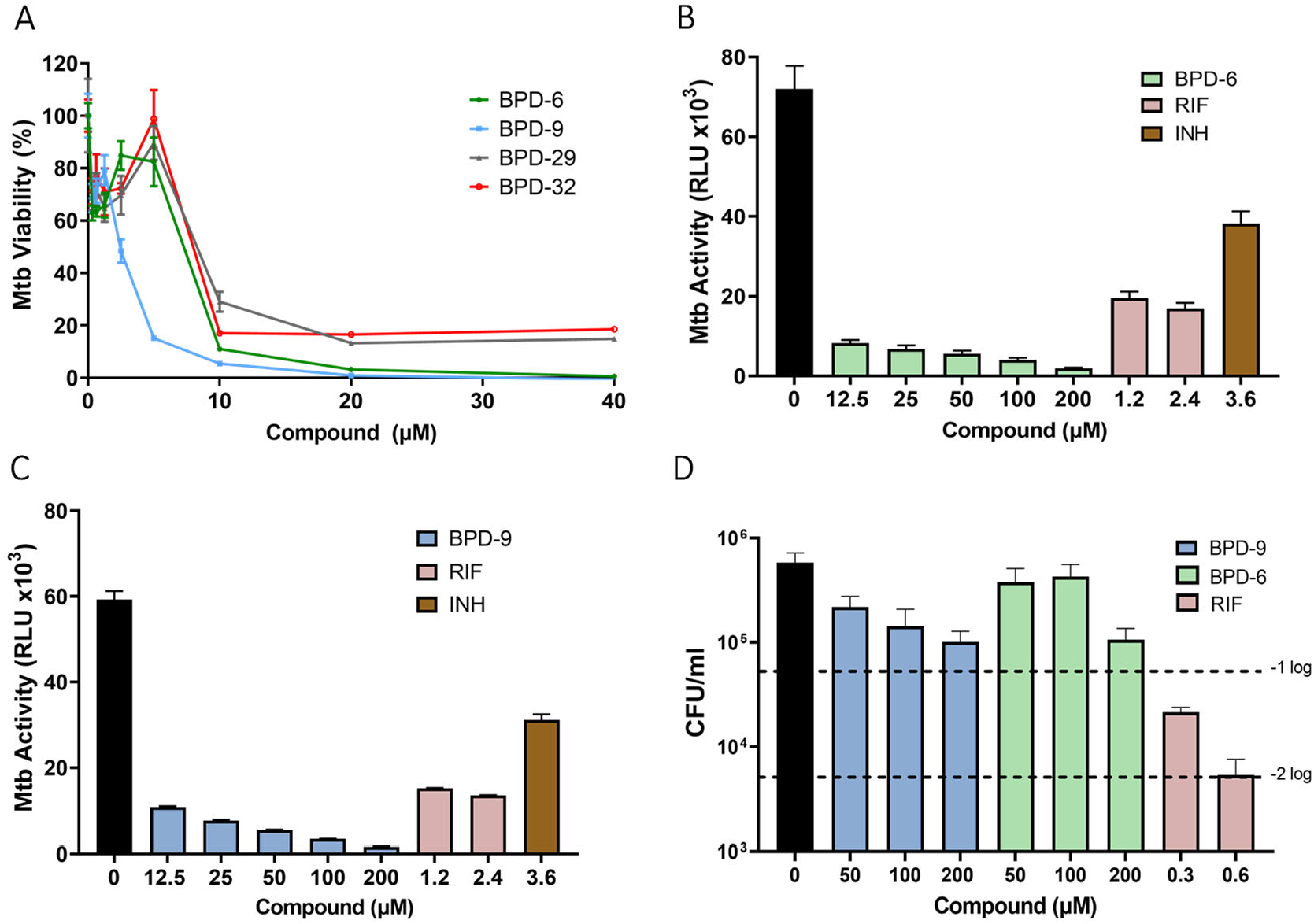
BPD-6 and BPD-9 significantly inhibit Mtb growth. (A) Dose-dependent activity of BPD-6, -9, -29, and -32 against Mtb mc^2^6206 was determined using the REMA assay. Mtb viability is normalized to maximal bacterial growth in the absence of compounds as 100%. (B, C) Mtb-*lux* was treated with BPD-6 (B) or BPD-9 (C) and RIF/INH as controls at the indicated concentrations for 20 h. The viability (activity) of Mtb-*lux* was determined by measuring the resulting luminescence of the strains (RLU, relative light unit). (D) Mtb mc^2^6206 were mock-treated or treated with BPD-6, BPD-9 and RIF for 20 h. Bacteria were serially diluted and select dilutions were inoculating on 7H10 agar plates for enumeration of colony forming units (CFU). Data represent the mean ± SEM of 3 independent replicates.

**Table 1.** MIC_90_ of BPD-6, BPD-9 and Sanguinarine against various bacteria species. n.a, no activity up to 100 μM; n.d., not determined. EC, *Escherichia coli*; LM, *Listeria monocytogenes*; PA, *Pseudomonas aeruginosa*; SE, *Salmonella enterica* Typhimurium.

| Compound<br>(μM) | *Mycobacterium* spp. | | | | Other bacterial species | | | | |
| --- | --- | --- | --- | --- | --- | --- | --- | --- | --- |
|  | Mtb<br>mc<sup>2</sup>6206 | *M. bovis*<br>BCG | *M.*<br>*kansasii* | *M.*<br>*smegmatis* | *EC* | *LM* | *PA01* | *PA14* | *SE* |
| BPD-6 | 10 | 2 | 12 | 3 | n.a | n.a | n.a | n.a | n.a |
| BPD-9 | 6 | 2 | 25 | 2 | n.a | n.a | n.a | n.a | n.a |
| Sanguinarine | 85 | 39 | > 40 | 38 | $\leq 24$<sup>17</sup> | n.d. | 99<sup>19</sup> | n.d. | n.d. |

To determine the efficacy of BPD-6 and -9 in the direct killing of Mtb compared to frontline TB drugs, including RIF and INH, we generated an auto-luminescent Mtb mc^2^6206 strain (Mtb*-lux*) to accelerate the measurement of bacterial viability upon treatment with BPD compounds. This recombinant Mtb*-lux* strain expresses a bacterial luciferase operon that allows the bacteria to generate a constitutive luminescence signal.^27,28^ Luminescence signal depends not only on the proteins of the *lux* operon, but also requires ATP and NADPH. As such, the relative luminescence unit (RLU) acts as a surrogate reporter for the viability of metabolically active Mtb. Treatment of Mtb*-lux* with BPD-6 and -9 for 24 hours resulted in a dose-dependent inhibition of Mtb viability (Figure 2B, C). Importantly, when used even at 1- to 2-fold the MIC_90_, both BPD-6 and -9 showed increased inhibition of Mtb viability compared to RIF or INH at 1.2 and 3.6 μM respectively, which are 10-fold their MIC_90_.^29^ These results indicate that the two compounds function rapidly to potently inhibit the metabolism and replication of Mtb. However, the use of luciferase as a readout for viability has limitations as bacteria with low metabolic activity may only produce non-detectable levels of luminescence signal. To assess whether the new BPD compounds possessed bactericidal ability, we performed the classical colony-forming unit (CFU) plating assay. Mtb treated with RIF at 5-fold its MIC_90_ showed the expected ∼2-log (99%) reduction in bacteria CFU (Figure 2D). However, both BPD-6 and -9 were unable to reach this level of efficacy even at 10-fold their MIC_90_ (Figure 2D). While BPD-6 and -9 can kill Mtb as shown by a near 1-log reduction in CFU at high concentrations > 100 μM (Figure 2D), it is clear that their bactericidal potency is limited. Instead, this suggests that BPD-6 and -9 act in a bacteriostatic manner. However, this does not discount the therapeutic potential of the compounds. For example, ethambutol, which acts through a bacteriostatic mechanism,^30^ is part of the regimen to treat active TB. As well, there is evidence that bacteriostatic macrolides have potential in the treatment of MDR-TB.^31–33^ Indeed, treatment of TB relies on multiple antibiotics to circumvent drug resistance and bacteriostatic compounds are often combined with bactericidal drugs to achieve the best effect.^34^ Given the bacteriostatic activity of BPD-6 and -9, attempts to generate escape mutants were unsuccessful. We also speculate that resistance to these compounds could be much slower to develop.

### BPD-6 and BPD-9 are active against non-replicating Mtb

Despite the lack of bactericidal activity, the BPD’s ability to inhibit the metabolic activity of Mtb is comparable to or better than RIF. Bacteriostatic drugs targeting Mtb are valuable if they can also inhibit NR-Mtb, which have increased tolerance to most antibiotics.^34^ To determine the efficacy of BPD-6 and BPD-9 against Mtb with low metabolic activity, we used the established low pH model to generate NR-Mtb.^7^ Mtb*-lux* was cultured in acidic phosphate citrate buffer without carbon or nitrogen supplements and the resulting NR-Mtb were used to assess the ability of the BPD compounds to inhibit their growth. The generated NR-Mtb showed very low levels of luciferase activity (Figure 3A), which supports that the bacteria are in a metabolically low, non-replicating state.^35,36^ NR-Mtb were incubated in complete media in the absence or presence of BPD compounds, or RIF and INH as controls for 20 hours. Untreated NR-Mtb recovered their ability to produce a luminescence signal, indicating their metabolic resuscitation and growth in this period of time (Figure 3B, C). In contrast, BPD-6 and -9-treated NR-Mtb were completely inhibited from metabolic resuscitation and growth, to levels comparable or even better than RIF used at a relatively high concentration (10-20x MIC_90_) (Figure 3B, C). While RIF possesses good inhibitory activity against NR-Mtb, INH only has limited efficacy against NR-Mtb *in vitro*.^7^ Indeed, in the same set of experiments, treatment of NR-Mtb with a high concentration of INH (10x MIC_90_) did not show any significant inhibition of growth (Figure 3B, C).

**Figure 3.**
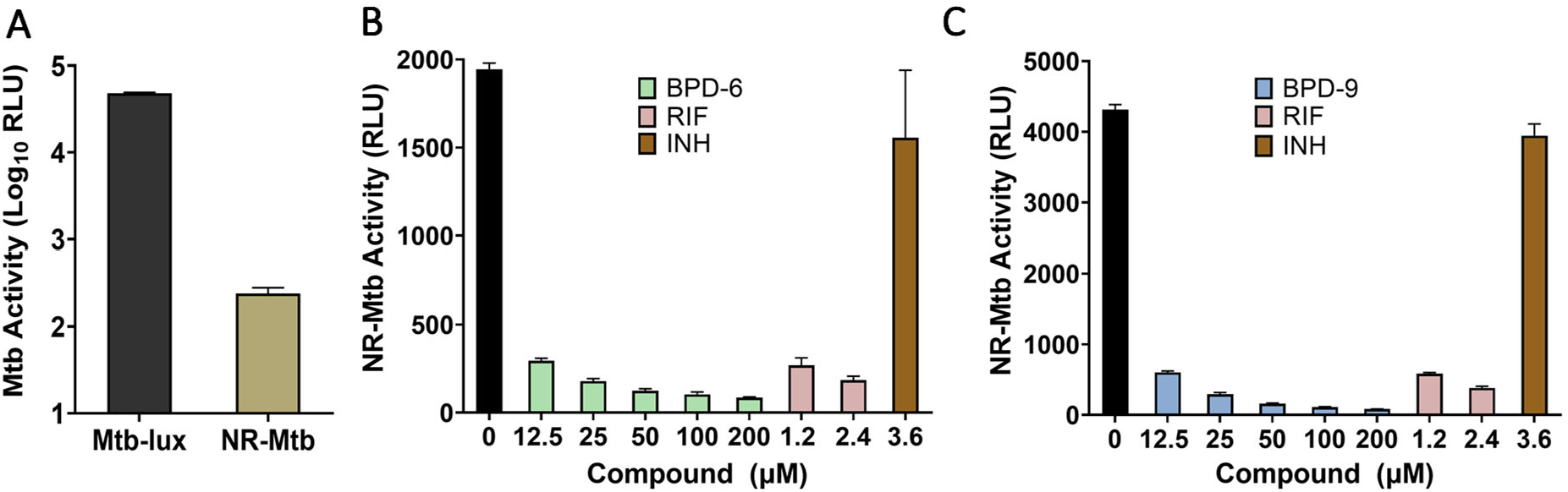
BPD-6 and BPD-9 are active against latent Mtb. (A) Log-phase Mtb-*lux* and non-replicating Mtb-*lux* (NR-Mtb) were resuspended to an OD_600_ of 0.03 in 7H9 media and their luminescence were measured immediately. (B, C) NR-Mtb were treated with BPD-6 (B) or BPD-9 (C), and RIF/INH as controls at the indicated concentrations for 20 hours. The activity of the compounds against NR-Mtb were then determined by measuring the resulting luminescence (RLU). Data represent the mean ± SEM of 3 independent replicates.

In physiological conditions, Mtb has the ability to maintain a dormant state within macrophages and reactivate upon perturbations to the immune system.^37^ Our *in vitro* model of NR-Mtb simulates this dormancy state, verified by the low level of luminescence (Figure 3A). This luminescence, controlled by the *lux* operon, exploits fatty aldehydes as substrates to produce fatty acids and light, meaning luminescent signal is dependent on an aerobic and metabolically active environment.^27^ In our experiments, we observed resuscitation of untreated NR-Mtb upon incubation in complete growth media, due to restoration of metabolic activity and fatty acid biosynthesis.^38^ However, BPD-treated bacteria remained in a low metabolic state. Considering that BPDs act as bacteriostatic agents, it is likely that their target and mechanism of inhibiting NR-Mtb are different than that of RIF. Many drug targets have been identified and well-studied in Mtb with low metabolic activity, though none are specifically involved in the process of reactivation.^6^ Given that the BPD compounds inhibit reactivation of NR-Mtb, the identification of a potentially novel drug target involved in the regulation of metabolic activity in NR-Mtb is an appealing topic for future research.

### BPDs exhibit reduced cytotoxicity and are effective against intracellular Mtb

The therapeutic potential of SG is limited given its cytotoxic properties due to non-specific targeting of essential eukaryotic proteins such as the Na^+^/K^+^-ATPase.^39^ Indeed, we show that human THP-1 macrophages treated with SG show significant cytotoxicity, with an IC_50_ of 10 μM (Figure 4A). The IC_50_ of SG is 4-fold lower than the MIC_50_ of SG against Mtb, rendering it useless for targeting intracellular Mtb, which is critical given that Mtb is an intracellular pathogen. In contrast, BPD-6 and BPD-9 showed a cytotoxicity IC_50_ of 40 μM and 15 μM in macrophages, respectively (Figure 4A). As such, BPD-9 and BPD-6 have a cytotoxic IC_50_ that is ∼5 to 6-fold higher than their respective MIC_50_ against Mtb. Taken together, the *in vitro* therapeutic ratio, defined as IC_50_/MIC_50_ for the respective compounds, illustrates the potential and improvement of BPD-6 and -9 compared to SG (Table 2). Despite their similar chemical structures, BPD-9 showed increased cytotoxicity compared to BPD-6, suggesting that a unique structural feature of BPD-9 contributes to its increased cytotoxicity. We speculate that minimizing the structural complexity of the BPD compounds will be key to reducing cytotoxicity while retaining activity.

**Table 2.** Therapeutic ratio (TR) of BPD-6, BPD-9, and SG.

| | Cytotoxicity<br>IC<sub>50</sub> (μM) | Mtb<br>MIC<sub>50</sub> (μM) | TR<br>(IC<sub>50</sub>/MIC<sub>50</sub>) |
| --- | --- | --- | --- |
| BPD-6 | 40.0 | 7.0 | 5.7 |
| BPD-9 | 15.0 | 3.0 | 5.0 |
| Sanguinarine | 10.0 | 40.0 | 0.25 |

**Figure 4.**
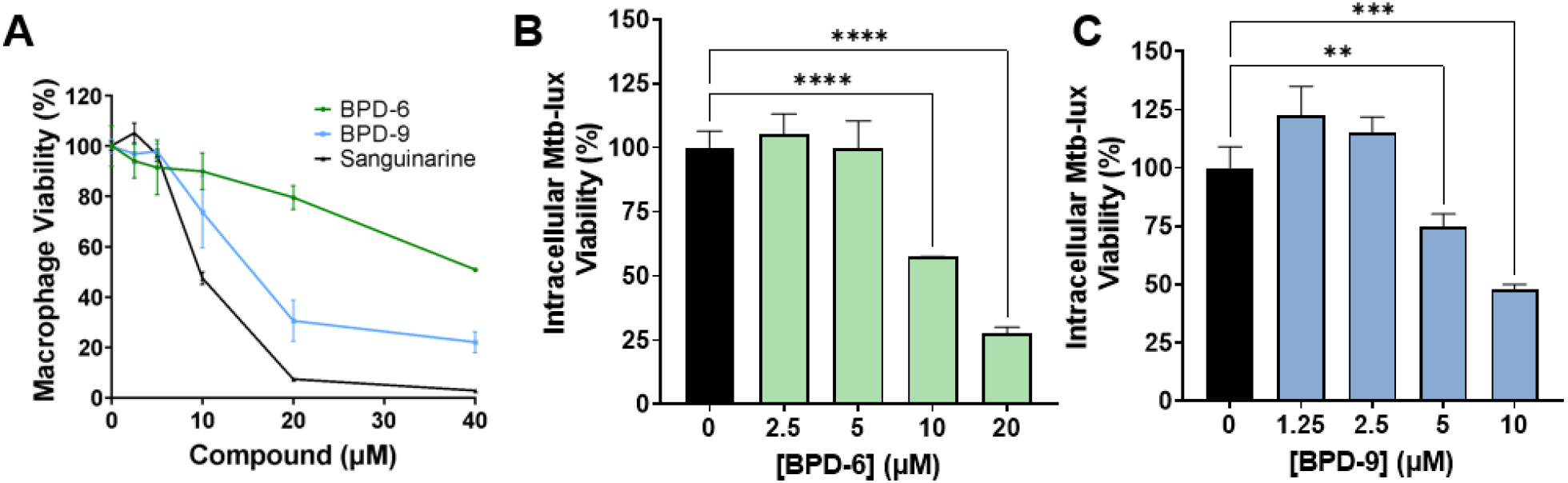
BPD-6 and BPD-9 inhibits intracellular Mtb. (A) The cytotoxicity of BPD-6, BPD-9, and Sanguinarine against THP-1 macrophages was determined at the indicated concentrations for 24 h. Macrophage viability was normalized relative to an untreated control as 100%. (B, C) THP-1 macrophages were infected with Mtb-*lux* at a multiplicity of infection (MOI) of 10. Infected cells were treated with BPD-6 (B) or BPD-9 (C) at the indicated concentrations for 24 h. The viability of intracellular Mtb was then determined by measuring the resulting luminescence (RLU). Data represent the mean ± SEM of 3 independent replicates. \*\**p* < 0.01; \*\*\**p* < 0.001; \*\*\*\**p* < 0.0001.

The reduced cytotoxicity of the BPD compounds enabled the evaluation of anti-Mtb efficacy in macrophage infection models. Mtb*-lux* has proven to be a reliable strain for determination of intracellular Mtb viability *in vitro* and *in vivo*.^7,27^ Treatment of infected macrophages with BPD-6 and BPD-9 at 10 μM (∼1x MIC_90_) effectively inhibited the intracellular growth of Mtb by 50% compared to untreated control cells (Figure 4B, C). BPD-9 showed better inhibition than BPD-6, starting at 5 μM. These results indicate that the BPD compounds remain effective at inhibiting Mtb replication inside the macrophage at near-MIC concentrations, which highlights the value of BPD-6 and -9 as novel compounds that have potential to treat TB.

### Combination treatment of BPD compounds with rifampicin improves anti-Mtb activity in axenic conditions and in infected macrophages

The combinatorial effectiveness of different antibiotics and drugs is one of the most emphasized aspects in drug development. To evaluate possible combinatorial effects of BPD-6 and BPD-9 with existing TB antibiotics, we adopted the combination checkerboard assay.^40^ This assay can be used to evaluate drug synergy through the calculation of the Fractional Inhibitory Concentration Index (FIC).^41^ Checkerboard experiments did not show synergistic effects between either BPD compound and RIF, INH, ethambutol (EMB), or moxifloxacin (MFX) as FICs were > 0.5 (data not shown). Importantly, there were no negative synergistic effects with any of the tested TB drugs. However, these assays revealed potential additive effects between BPD-6 and -9 with RIF. To confirm these observations, we performed REMA assays with Mtb treated either with BPD-6, -9, or RIF alone, or in combination at sub-MIC concentrations. These assays showed that the combined treatment of Mtb with BPD-6 or BPD-9 with RIF significantly decreased Mtb growth compared to single treatments (Figure 5A, B). The combination effect was specific to RIF, and did not occur in combination with INH, EMB, and MFX. Importantly, the increased activity of BPD-6 and -9 in combination with RIF translated to intracellular conditions, where we observed a decrease in Mtb viability in macrophages treated with the combination of BPD-6 or -9 and RIF compared to single treatments (Figure 5C, D). Given that the combination effect with BPDs only occurred with RIF, we speculate that the mechanism of action for BPDs is distinct from RIF. Combination effects are desirable since it can lower the effective concentrations of each drug to minimize cytotoxicity.

**Figure 5.**
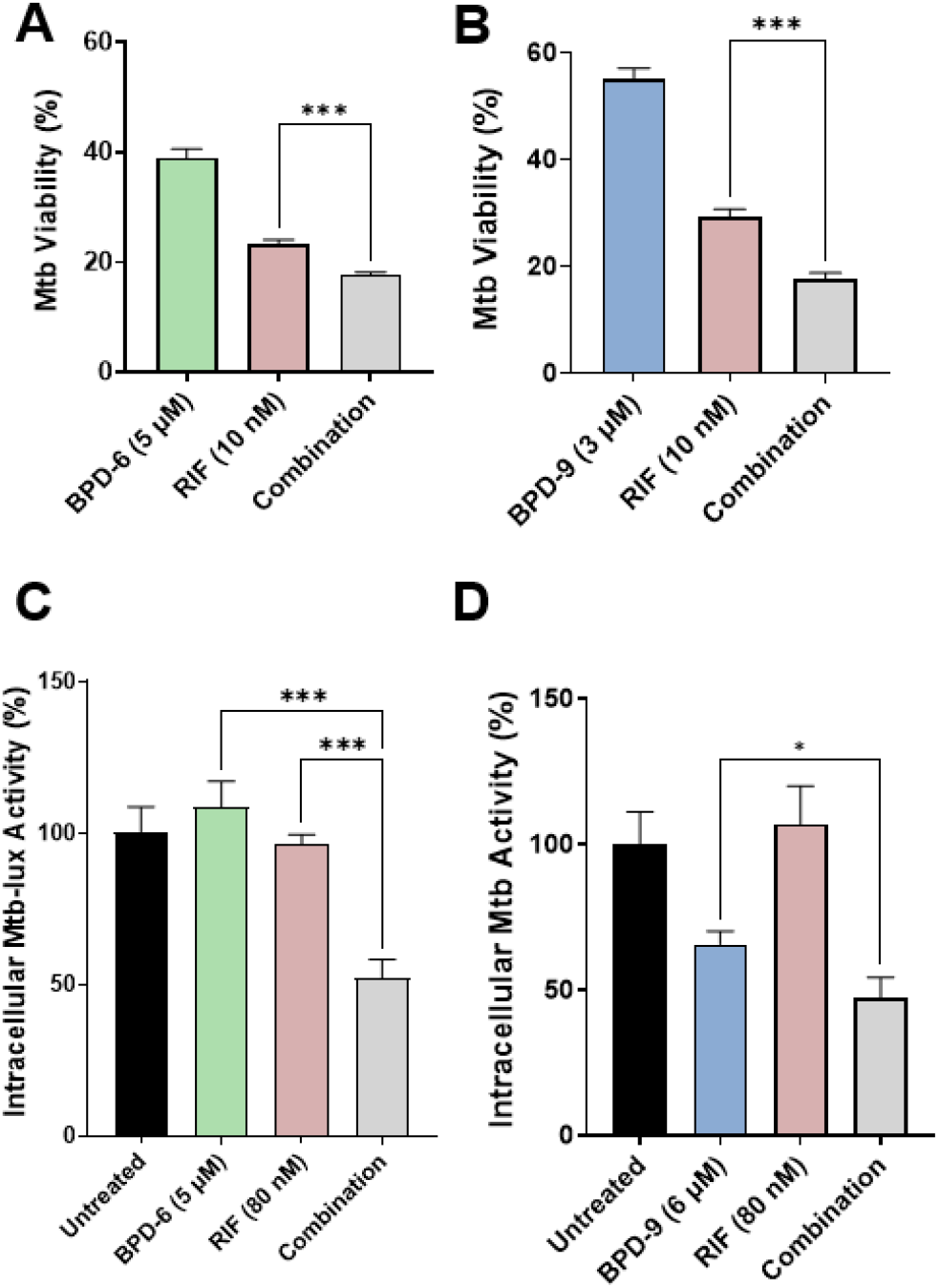
BPD-6 and BPD-9 combines with RIF to improve anti-Mtb activity. (A, B) Activity of BPD-6 (A) or BPD-9 (B) alone or in combination with RIF against Mtb mc^2^6206 was determined using the REMA assay. Mtb viability is normalized to maximal bacterial growth in the absence of compounds as 100%. (C, D) THP-1 macrophages were infected with Mtb-*lux* and treated with BPD-6 (C) or BPD-9 (D) alone or in combination with RIF for 24 h. Intracellular Mtb viability was determined by measuring the resulting luminescence. Data represent the mean ± SEM of 3 independent replicates. \**p* < 0.05; \*\*\**p* < 0.001.

### BPD-6 and BPD-9 accumulate within the bacterial cell

. To gain insight into how the BPD compounds may exert their antibacterial functions, we sought to investigate the dynamics of the compounds’ interaction with bacteria. Interestingly, BPD-6 and BPD-9 have fluorescence properties (peak excitation/emission wavelengths of 420/485 nm) that can be exploited for visualization and quantification (Figure S3). To examine the uptake and localization of the compounds inside the bacteria, Mtb-RFP^42^ treated with sub-MIC concentrations of BPD-6 and BPD-9 were visualized by epifluorescence microscopy. Fluorescence of BPD-6 and BPD-9 co-localized with RFP signal and was evenly distributed throughout the bacteria, indicating a possible accumulation of the compounds in the cytosol of Mtb cells (Figure 6A). However, a portion of the Mtb did not appear to accumulate the compounds, suggesting incomplete uptake at this specific concentration.

**Figure 6.**
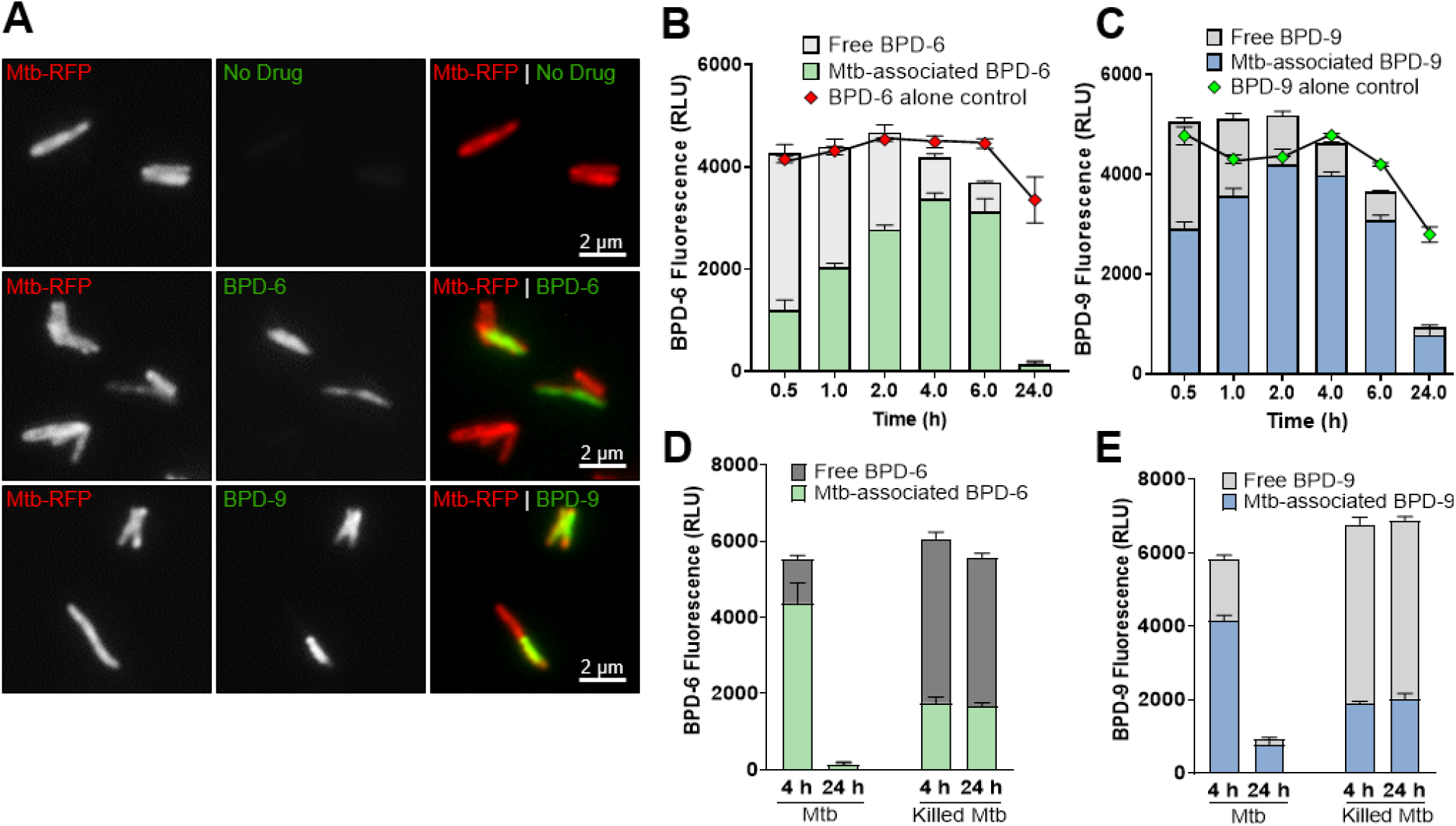
BPD-6 and BPD-9 accumulate within Mtb cells. (A) Representative fluorescence image of Mtb-RFP treated with 5 μM BPD-6 or BPD-9 for 4 h. (B, C) Mtb mc^2^6206 were treated with 5 μM BPD-6 (B) or BPD-9 (C) for the indicated time points. The resulting fluorescence of the BPD compounds in the Mtb pellet (Mtb-associated BPD) and the supernatant (free BPD) were measured. A “BPD-alone control” in the absence of Mtb was included to measure the endogenous decay of BPD’s fluorescence. (D, E) Live or gentamicin-killed Mtb mc^2^6206 were treated with 5 μM BPD-6 (D) or BPD-9 (E) for 4 h and 24 h, and the accumulation of the compounds in Mtb was measured by the resulting fluorescence. Data represent the mean ± SEM of 3 independent replicates.

To quantify the accumulation of the compounds bound to or inside the bacteria over a period of 24 h, we measured fluorescence in the Mtb pellet (Mtb-associated) and the supernatant (free), which would contain compound that was either not taken up or that was effluxed by the bacteria. Maximal uptake of BPD-6 and BPD-9 occurred within 4 hours of treatment, which corresponded to a relatively low percent of unassociated compound remaining in the supernatant (Figure 6B, C). The relatively quick time for the BPD compounds to accumulate within the Mtb is consistent with the chemical properties of benzo[c]phenanthridine compounds, which tend to be nonpolar with a high level of molecular planarity, enabling them to more easily enter the bacterial cell.^43^ Interestingly, we observed a dramatic decrease in fluorescence in the Mtb pellet at 24 h post-treatment (Figure 6B, C). The magnitude of this decrease in fluorescence is significantly higher than the natural decrease observed when the compounds are incubated in the exact same conditions without Mtb. There was also no evidence of increased efflux in the supernatant. A possible explanation for the decrease in fluorescence of the compounds following prolonged incubation with Mtb is that the bacteria is able to metabolise the compound, or that interaction with its cognate target within Mtb interferes with the intrinsic fluorescence of the compound. Consistent with the hypothesis of compound metabolizing, we show that only treatment of live, but not killed Mtb (Figure S4), results in a decrease in fluorescence at 24 h post-treatment (Figure 6D, E). In addition, metabolically active live bacteria are required for uptake of the compound as killed bacteria accumulate significantly lower amounts of both BPD-6 and BPD-9 after 4 h of treatment (Figure 6D, E).

### BPD-6 and BPD-9 are effective against virulent and clinical MDR Mtb strains

Given the increasing prevalence of MDR-TB, novel anti-Mtb compounds that retain their efficacy against clinical and MDR strains of Mtb are highly desirable. We show that both BPD-6 and BPD-9 retain their efficacy in the low micromolar MIC range against both virulent laboratory strains (Mtb H37Rv and Mtb Erdman) and the hypervirulent Mtb HN878 strain (Figure 7A-C and Table 3). Importantly, we examine the efficacy of both compounds against a panel of 5 clinical drug resistant Mtb isolates (50: INH^R^; 105: PZA^R^, INH^R^, STP^R^; 116: INH^R^, RIF^R^, STP^R^; 151: PZA^R^, INH^R^, RIF^R^, STP^R^; 217: PZA^R^, INH^R^, RIF^R^, ETB^R^). Strikingly, both BPD-6 and BPD-9 effectively inhibited the growth of all 5 clinical MDR Mtb strains in the low micromolar range, whereas resistance to rifampicin and isoniazid was observed in the corresponding isolates (Figure 7D-H and Table 3). Collectively, our data show that BPDs are effective against multiple strains of virulent and MDR Mtb and that their mechanism of action is distinct from that of the frontline TB drugs, including rifampicin, isoniazid, pyrazinamide, streptomycin, and ethambutol.

**Figure 7.**
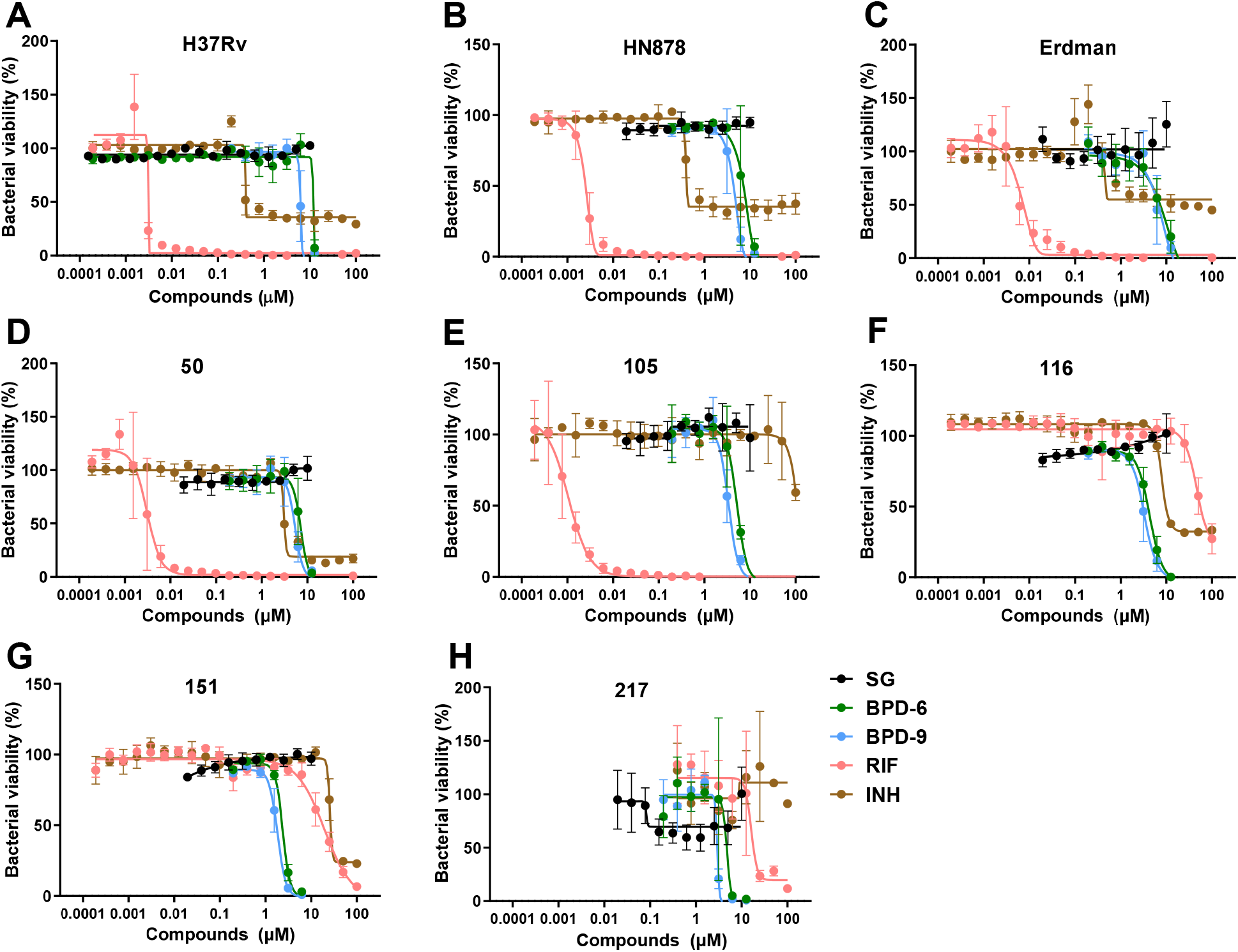
BPD-6 and BPD-9 are active against virulent and clinical Mtb strains. Dose-dependent activity of sanguinarine (SG), BPD-6, BPD-9, rifampicin (RIF), and isoniazid (INH) against the Mtb strains: H37Rv (A), HN878 (B), and Erdman (C), clinical isolate #50 (D), #105 (E), #116 (F), #151 (G), and #217 (H) was determined using the REMA assay. Mtb viability is normalized to maximal bacterial growth in the absence of compounds as 100%. Data represent the mean ± SEM of 4 replicates.

**Table 3.**
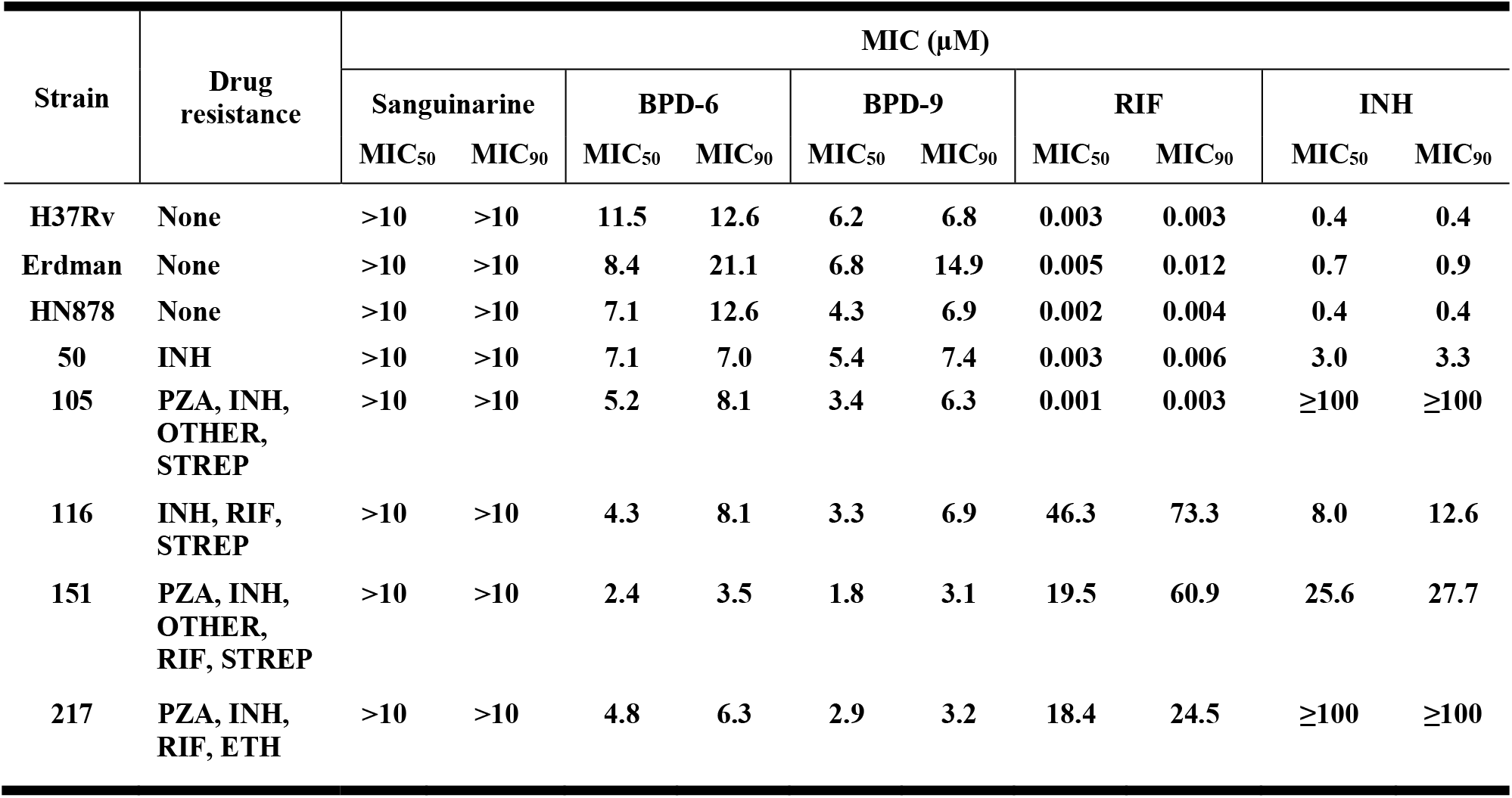
MIC of sanguinarine, BPD-6, BPD-9, rifampicin, and isoniazid against various laboratory strains and clinical isolates of *M. tuberculosis*. RIF, rifampicin; INH, isoniazid; PZA, pyrazinamide; STREP, streptomycin; ETH, ethambutol.

## CONCLUSIONS

In summary, we report the design and synthesis of a group of benzo[c]phenanthridine compounds derived from sanguinarine that possesses potent and specific antimycobacterial activity. Our data also provide evidence for the active pharmacophore of benzo[c]phenanthridine compounds and the moieties that impact cytotoxicity and activity. We provide biological evidence that these compounds inhibit metabolically inactive non-replicating Mtb, intracellular Mtb, and multiple clinical MDR-TB strains in the low micromolar range. The potency, specificity and cytotoxicity are all superior relative to sanguinarine, and suggests a unique mode of action that is distinct from all frontline TB antibiotics. The selective ability to inhibit multiple species of *Mycobacterium* also supports a target that is unique to this genus, which will be valuable for the development of more potent antibiotics that specifically target mycobacteria.

## EXPERIMENTAL SECTION

### Chemistry

#### General Procedures

Starting materials, reagents, and solvents were purchased from commercial suppliers and used without further purification. Silica gel for column chromatography was of 200-300 mesh particle size, EA/PE mixture and DCM/MeOH mixture were used for purification. NMR spectra were recorded at room temperature on Bruker Avance III 400 Spectrometer (400 MHz). Chemical shifts are given in ppm and coupling constants in Hz. ^1^H spectra were calibrated in relation to the reference measurement of TMS (0.00 ppm). ^13^C spectra were calibrated in relation to deuterated solvents. High-resolution mass spectra (HRMS) were performed using the Agilent G6520 Q-TOF high-resolution mass spectrometer. The purity of active compounds BPD-6, -9, -19, -29, -32 were determined by high performance liquid chromatography (HPLC). HPLC conditions: Agilent 1260 with a XBridgeTM C18 column (4.6 mm × 150 mm, 5 μm); T = 40 °C; flow rate = 0.5 mL/min. All final compounds were confirmed to have a purity of over 95%.

#### Synthetic route for BPD-6 and BPD-9 (Scheme 1)

For the synthesis of the most potent compounds BPD-6 and BPD-9, an F-C acetylation was utilized starting from anisole and veratrole respectively. Treatment anisole or veratrole with succinic anhydride under AlCl_3_ condition provided **A2** after work up. Next, **A2** was converted into **A4** by reduction with Et_3_SiH and dehydration in PPA and aromatization was conducted by subsequently treating **A4** with Br_2_ and DBU. The corresponding arylamine **A9** was prepared by S_N_2, Smiles rearrangement and hydrolysis reactions. Finally, **A9** was treated with formyl acetate, MeI, followed by a Suzuki coupling with aminophenyl borate and Vilsmeier-Haack reaction in POCl_3_ condition for ring closure to yield BPD-6 and BPD-9. The detailed synthetic procedure are as follows:

**Step a**: To a mixture of **A1** (1.0 equiv) and succinic anhydride (1.25 equiv) in nitrobenzene (2.67 M), AlCl_3_ (2.0 equiv) was added in ice bath, and the reaction mixture was stirred at room temperature and monitored by TLC. After completion of the reaction, the reaction was quenched by ice water, and washed with water and EA. The organic phase was dried over Na_2_SO_4_ and concentrated under reduced pressure. The residue was recrystallized to afford **A2**.

**Step b and c**: A mixture of **A2** (1.0 equiv) and Et_3_SiH (2.5 equiv) in TFA (1.4 M) was stirred at 100 °C for 7 h. The reaction mixture was cooled to room temperature and then diluted with water. The product was extracted with EA, then the organic extract was dried over Na_2_SO_4_ and concentrated under reduced pressure. Polyphosphoric acid (0.5 mL/mmol) was added to the crude product dropwise at room temperature, and the reaction mass was heated at 80 °C and monitored by TLC. After the completion of reaction, the mixture was diluted with EA in 0 °C and quenched by ice water. The product was extracted with EA, and the combined organic phase was dried over Na_2_SO_4_ and concentrated under reduced pressure. The residue was purified by silica gel chromatography to afford **A4**.

**Step d:** A mixture of **A4-2** in 48% HBr (0.325 M) was refluxed at 125 °C for 6 h. The reaction was cooled to room temperature, the brown crystal was filtered and dried to afford the crude product. The brown crystal was then diluted in DMF (0.2 M), K_2_CO_3_ (5.0 equiv) and CH_2_Br_2_ (1.2 equiv) were added. The mixture was heated at 85 °C for 2 h. After the reaction was cooled to room temperature, the solvent was removed and the residue was diluted with EA. The solution was washed with water, dried over Na_2_SO_4_, and purified by silica gel chromatography to afford **A4-3**.

**Step e and f:** To a mixture of **A4** in chloroform (1.5 M), Br_2_ was added dropwise at room temperature, and the reaction was stirred at room temperature and monitored by TLC. After the completion of reaction, the mixture was quenched by aqueous Na_2_SO_3_, washed with water and extracted with EA. The organic phase was dried over Na_2_SO_4_ and concentrated under reduced pressure. The crude product was dissolved in MeCN (1.0 M), and DBU (1.5 equiv) was added at 50 °C and the mixture was then stirred at 50 °C for 10 min. The reaction was diluted with EA, washed with water, dried over Na_2_SO_4_ and concentrated under reduced pressure, then purified by silica gel chromatography to give **A6**.

**Step g:** To a solution of **A6** (1.0 equiv) in DMF (0.2 M), NaOH (2.0 equiv) was added. After stirring at room temperature for 30 min, 2-bromo-2-methylpropanamide (2.0 equiv) was added, and then stirred for 2 h at ambient temperature. The mixture was diluted with EA, washed with water and dried over Na_2_SO_4_, then concentrated under reduced pressure and purified by silica gel chromatography to give **A7**.

**Step h and i**: To a solution of **A7** in DMF/DMTP (4:1, 0.22 M), NaH (2.0 equiv) was added at room temperature, and the mixture was heated at 100 °C for 4 h. The reaction was cooled to room temperature and quenched by aqueous NH_4_Cl. The product was extracted with EA, dried over Na_2_SO_4_, concentrated under reduced pressure, and purified by silica gel chromatography to give **A8. A8** was then dissolved in MeOH/H_2_O (1:1, 0.15 M), and NaOH (w/w 80%) was added. The resulting mixture was then heated at 100 °C and monitored by TLC. After completion of the reaction, the mixture was washed with water, extracted with EA, concentrated under reduced pressure and purified by gel chromatography to afford **A9**.

**Step j and k:** To a solution of **A9** (1.0 equiv) in THF (0.32 M), formyl acetate (3.0 equiv) was added and the mixture was stirred for 30 min at room temperature. The product was washed with water and extracted with EA. The organic phase was dried over Na_2_SO_4_ and concentrated under reduced pressure to afford **A10**. A mixture of **A10** (1.0 equiv) and NaH (2.0 equiv) in DMF (0.27 M) was stirred for 10 min at room temperature, then MeI (3.0 equiv) was added and the reaction was stirred for 1 h at ambient temperature. After completion of the reaction, the mixture was washed with water and extracted with EA, dried over Na_2_SO_4_, and concentrated under reduced pressure to afford **A11**.

**Step l and m:** To a solution of **A11** (1.0 equiv) in 1,4-dioxane/H_2_O (1:1, 0.075 M), N,N-dimethyl-3- (4,4,5,5-tetramethyl-1,3,2-dioxaborolan-2-yl)aniline (1.5 equiv), Pd(PPh_3_)_4_ (5 mol%) and K_3_PO_4_ (1.5 equiv) were added under nitrogen atmosphere, and the suspension was heated at 90 °C for 3 h. The reaction was cooled to room temperature, washed with water and extracted with EA, then concentrated under reduced pressure to give a yellow oil. Next, POCl_3_ was added (1.0 mL/mmol) to the crude product and the mixture was heated at 60 °C and monitored by TLC. After completion of the reaction, the mixture was diluted with DCM, then yellow solid appeared and was filtered to afford the product **BPD-6** and **BPD-9**. Total yield for BPD-6 was 2.6% and 1.2% for BPD-9.

### Biology

#### Cell culture

THP-1 monocytes (ATCC TIB-202) were maintained at 37°C in a humidified atmosphere of 5% CO_2_ in RPMI 1640 media (Gibco, Gaithersburg, MD) supplemented with 10% heat-inactivated fetal bovine serum (FBS), 10 mM HEPES, penicillin (100 I.U./ml), streptomycin (100 μg/ml), and 2 mM L-glutamine purchased from Gibco. For differentiation into THP-1 macrophages, cells were resuspended in complete RPMI 1640 media (without antibiotics) and incubated with 100 ng/mL phorbol 12-myristate 13-acetate (PMA, Alfa Aesar, Haverhill, MA). Cells were seeded at 50,000 cells/well into 96-well plates and incubated at 37°C for 3 days.

#### Bacterial culture

The *Mycobacterium tuberculosis* H37Rv-derived auxotrophic strain mc^2^6206 (Δ*PanCD*Δ*LeuCD*)^44^, virulent strains (Mtb Erdman, H37Rv, HN878), clinical MDR strains (isolates #50, 105, 116, 151, 217), *Mycobacterium kansasii* (ATCC 12478), and *Mycobacterium bovis* BCG (BCG Pasteur) were cultured in Middlebrook 7H9 media (BD Biosciences, Franklin Lakes, NJ) supplemented with 0.05% Tween 80 (Acros Organics, Fair Lawn, NJ), 0.2% glycerol (Fisher Chemical, Waltham, MA), and 10% OADC (BD Biosciences). For the Mtb mc^2^6206 strain, D-pantothenic acid (24 μg/ml, Alfa Aesar) and L-leucine (50 μg/ml, Alfa Aesar) were also added. For *M. kansasii*, 5 μg/ml streptomycin was also added (Fisher Scientific, Waltham, MA). *Mycobacterium smegmatis* (ATCC 700084) was maintained in 7H9 media supplemented with 10% ADS enrichment: 50 g bovine serum albumin (VWR, Radnor, PA), 20 g dextrose (Fisher Scientific), 0.85% (w/v) sodium chloride (Fisher Chemical), 0.2% glycerol, and 0.05% tyloxapol (Acros Organics). Clinical MDR Mtb isolates were obtained from the McGill International TB Centre (Montreal, Canada).

The recombinant bioluminescent strain Mtb*-lux* was generated by transforming the pMV306hsp+LuxG13 plasmid into Mtb-mc^2^6206.^27,28^ pMV306hsp+LuxG13 was a gift from Brian Robertson & Siouxsie Wiles (RRID: Addgene_26161). This plasmid encodes a bacterial luciferase-expressing operon *lux* and G13 promoter, allowing the constant production of luminescence signal by metabolically active bacteria. The strain was maintained in complete 7H9 media supplemented with 30 μg/ml kanamycin (Fisher Scientific). Mtb mc^2^6206 expressing tdTomato (Mtb-RFP) was generated previously.^42^

*Escherichia coli* strain NEB Stable (New England Biolabs, Ipswich, MA), *Pseudomonas aeruginosa* strains PA01 and PA14 (generous gift from Dr. Thien-Fah Mah, University of Ottawa), *Salmonella enterica* serovar Typhimurium strain SL1344 and *Listeria monocytogenes* strain 10403s (both generous gifts from Dr. Subash Sad, University of Ottawa) were cultured in Lysogeny Broth (LB, Fisher Scientific). Bacteria were inoculated into LB from frozen glycerol stocks and grown at 37°C with shaking (200 rpm).

#### Resazurin microtiter assay (REMA)

Bacteria were diluted to an initial OD of 0.02 at mid-log stage and incubated with sanguinarine (Tocris Bioscience, Toronto, Canada), RIF (Fisher Scientific), INH (Acros Organics), EMB (Alfa Aesar), MFX (Alfa Aesar), or BPD compounds at 37°C. The incubation time was 5 days for Mtb and BCG, 20 h for *M. smegmatis* and 3 days for *M. kansasii*. Then, resazurin (Sigma-Aldrich, St. Louis, MO) solution was added to the wells to achieve a final concentration of 100 μM resazurin and 0.5% Tween 80. The mixtures were incubated at 37°C before measurement of fluorescence with the Synergy H1 Microplate Reader (BioTek, Winooski, VT). The readout was performed after 6 hours for Mtb, *M. kansasii*, and *M. bovis*, and 1 hour for *M. smegmatis*. For THP-1 macrophages, the same concentration of resazurin was added into the wells and fluorescence signal was recorded 2 hours after incubation.

#### Mtb viability assays

##### Luminescence

The direct killing ability of the compounds were assessed by the luminescence signal readout of Mtb*-lux*. Mid-log Mtb*-lux* was diluted to an OD of 0.03 and added into 96-well white plates at 100 μl in each well. 100 μl of serial diluted BPD-6 and 9 (200-25 μM), RIF (1.2 μM, Fisher Scientific) and INH (3.6 μM, Acros Organics) were added into the wells. A readout of luminescence signal was performed immediately after plate setup (time 0). The plates were incubated in 37°C without shaking for 20 hours, followed by a final readout. Luminescence was measured with the Synergy H1 Microplate Reader with an optimized integration time of 10 seconds.

##### Colony forming unit (CFU) plating

Middlebrook 7H10 agar plates (BD Biosciences) supplemented with 0.5% glycerol, 10% OADC, 24 μg/ml D-pantothenic acid, and 50 μg/ml L-leucine were prepared. Mid-log Mtb was washed and treated with serial diluted compounds (200-50 μM), RIF (250 ng/ml, 500 ng/ml) and untreated control in 96-well plates. The plates were incubated at 37°C for 20 hours. The bacteria in each well were serial diluted by 10-fold. The last four dilutions (10^−2^, 10^−3^, 10^−4^, 10^−5^) were inoculated on 7H10 agar plates. After 3 weeks of incubation at 37°C, each plate was counted for colonies and calculated for correlative CFU/ml.

#### Intracellular Mtb survival assay

THP-1 cells were differentiated into macrophages as described above. The amount of Mtb*-lux* was counted for a multiplicity of infection (MOI) of 10 using the conversion of 3×10^8^ bacteria/ml for OD 1.0. Log-phase bacteria were resuspended in RPMI 1640 infection media: supplemented with 10% human serum (Millipore Sigma, Burlington, MA), 10 mM HEPES, 2 mM glutamine, D-pantothenic acid (24 μg/ml) and L-leucine (50 μg/ml). After differentiation, the THP-1 derived macrophages were washed with PBS and recovered in infection media for 1 hour, and then infected with Mtb*-lux* for 4 hours. Extracellular Mtb*-lux* was removed by three PBS washes, followed by addition of compounds. Intracellular survival of Mtb*-lux* was quantified 24 h post-infection by luminescence readouts with the Synergy H1 Microplate Reader with an integration time of 10 seconds.

#### Generation of non-replicating Mtb (NR-Mtb)

Non-replicating Mtb (NR-Mtb) was generated using an established low-pH model.^7^ Mid-log Mtb*-lux* was washed with PBS (1X) and transferred into PCB media: 1X phosphate-citrate buffer (PCB, composed of 0.2 M sodium phosphate (Fisher Scientific) with 0.1 M citric acid (Fisher Scientific) at pH 4.5) supplemented with 0.05% tyloxapol. The OD was adjusted to 0.5. Bacteria were maintained at 37°C for 7 days. OD was measured every day to confirm the non-growing state. For treatments with compounds, non-replicating Mtb*-lux* was removed from PCB buffer on day 7 and resuspended in nutritional 7H9 media directly before compound treatments. RLU viability readouts were measured as described above.

#### Measurement of BPD accumulation within Mtb

Log-phase bacteria were washed and resuspended with PBS to OD 0.1 in Eppendorf tubes, and then treated with the compounds at the indicated concentrations at 37°C. At various time points, the bacteria-compound suspension was centrifuged at 8,000 x *g* for 3 minutes. Supernatants were transferred into 96-well plates, while the remaining pellets were washed 2 times with PBS. After the last PBS wash, pellets were resuspended and transferred into 96-well plates. Fluorescence was measured at an excitation/emission of 420/485 nm using the Synergy H1 Microplate Reader. Mtb-*lux* were treated with 50 μg/ml gentamicin (Calbiochem, San Diego, CA) for 24 h to generate killed Mtb. The RLU of killed Mtb were measured to confirm death of the bacteria.

#### Fluorescence microscopy

Mtb-RFP were treated with 5 μM BPD-6, BPD-9, or solvent for 4 hours. Bacteria were washed with PBS (1X, supplemented with 0.05% Tween 80) after incubation and fixed with 4% formaldehyde for 15 minutes. Slides were dried at 37°C and mounted with FluoroSave Reagent (Calbiochem). Samples were imaged with Invitrogen EVOS FL Auto Imaging System using 100X objective. Fluorescent filters were built-in GFP filter (excitation/emission: 470(22)/525(50) nm) and RFP filter (excitation/emission: 530(40)/593(40) nm). Images were analyzed with ImageJ software.

#### Statistical analysis

Statistical analysis was performed using the unpaired Student’s t-test or one-side ANOVA on GraphPad Prism Version 9. Values of *p* < 0.05 were considered as statistically significant.

## Associated Content

Supporting Information: *Supplementary Figures S1-4*.

## Author contributions

Z.S., YC.L., A.L., M.A.B., and J.S. conceived and designed experiments, and analyzed data; L.C., Z.X., and W.Y. designed and synthesized the chemical compounds; Z.S., YC.L., A.L., and S.B. performed experiments; Z.S., YC.L., W.Y., M.A.B., and J.S. wrote and edited the manuscript. All authors read and approved the final manuscript. The authors declare no competing financial interest

## Acknowledgments

We thank the support staff at the Cell Biology and Image Acquisition Core (CBIA) and Common Equipment and Technical Service (CETS) at the University of Ottawa for their expert technical assistance. We thank Dr. Thien-Fah Mah and Dr. Subash Sad at the University of Ottawa for generous gift of bacterial strains used in this study.

## References

(1) Chakaya, J.; Khan, M.; Ntoumi, F.; Aklillu, E.; Fatima, R.; Mwaba, P.; Kapata, N.; Mfinanga, S.; Hasnain, S. E.; Katoto, P. D. M. C.; Bulabula, A. N. H.; Sam-Agudu, N. A.; Nachega, J. B.; Tiberi, S.; McHugh, T. D.; Abubakar, I.; Zumla, A. Global Tuberculosis Report 2020 - Reflections on the Global TB Burden, Treatment and Prevention Efforts. Int. J. Infect. Dis. 2021, S1201-9712(21)00193-4.

(2) Zuniga, E. S.; Early, J.; Parish, T. The Future for Early-Stage Tuberculosis Drug Discovery. Future Microbiol. 2015, 10 (2), 217–229.

(3) Kumar, A.; Chettiar, S.; Parish, T. Current Challenges in Drug Discovery for Tuberculosis. Expert Opin. Drug Discov. 2017, 12 (1), 1–4.

(4) Ehrt, S.; Schnappinger, D.; Rhee, K. Y. Metabolic Principles of Persistence and Pathogenicity in Mycobacterium Tuberculosis. Nat. Rev. Microbiol. 2018, 16 (8), 496–507.

(5) Batyrshina, Y. R.; Schwartz, Y. S. Modeling of Mycobacterium Tuberculosis Dormancy in Bacterial Cultures. Tuberc. Edinb. Scotl. 2019, 117, 7–17.

(6) Gupta, V. K.; Kumar, M. M.; Singh, D.; Bisht, D.; Sharma, S. Drug Targets in Dormant Mycobacterium Tuberculosis: Can the Conquest against Tuberculosis Become a Reality? Infect. Dis. Lond. Engl. 2018, 50 (2), 81–94.

(7) Early, J. V.; Mullen, S.; Parish, T. A Rapid, Low PH, Nutrient Stress, Assay to Determine the Bactericidal Activity of Compounds against Non-Replicating Mycobacterium Tuberculosis. PloS One 2019, 14 (10), e0222970.

(8) Njire, M.; Tan, Y.; Mugweru, J.; Wang, C.; Guo, J.; Yew, W.; Tan, S.; Zhang, T. Pyrazinamide Resistance in Mycobacterium Tuberculosis: Review and Update. Adv. Med. Sci. 2016, 61 (1), 63–71.

(9) Mahajan, R. Bedaquiline: First FDA-Approved Tuberculosis Drug in 40 Years. Int. J. Appl. Basic Med. Res. 2013, 3 (1), 1–2.

(10) Nasto, B. Not-for-Profit to Launch Antibiotic against Drug-Resistant Tuberculosis. Nat. Biotechnol. 2019.

(11) Mdluli, K.; Kaneko, T.; Upton, A. Tuberculosis Drug Discovery and Emerging Targets. Ann. N. Y. Acad. Sci. 2014, 1323, 56–75.

(12) Atanasov, A. G.; Zotchev, S. B.; Dirsch, V. M.; International Natural Product Sciences Taskforce; Supuran, C. T. Natural Products in Drug Discovery: Advances and Opportunities. Nat. Rev. Drug Discov. 2021, 20 (3), 200–216.

(13) Sensi, P. History of the Development of Rifampin. Rev. Infect. Dis. 1983, 5 Suppl 3, S402–406.

(14) Dashti, Y.; Grkovic, T.; Quinn, R. J. Predicting Natural Product Value, an Exploration of Anti-TB Drug Space. Nat. Prod. Rep. 2014, 31 (8), 990–998.

(15) Grenby, T. H. The Use of Sanguinarine in Mouthwashes and Toothpaste Compared with Some Other Antimicrobial Agents. Br. Dent. J. 1995, 178 (7), 254–258.

(16) Tenenbaum, H.; Dahan, M.; Soell, M. Effectiveness of a Sanguinarine Regimen after Scaling and Root Planing. J. Periodontol. 1999, 70 (3), 307–311.

(17) Hamoud, R.; Reichling, J.; Wink, M. Synergistic Antibacterial Activity of the Combination of the Alkaloid Sanguinarine with EDTA and the Antibiotic Streptomycin against Multidrug Resistant Bacteria. J. Pharm. Pharmacol. 2015, 67 (2), 264–273.

(18) Beuria, T. K.; Santra, M. K.; Panda, D. Sanguinarine Blocks Cytokinesis in Bacteria by Inhibiting FtsZ Assembly and Bundling. Biochemistry 2005, 44 (50), 16584–16593.

(19) Falchi, F. A.; Borlotti, G.; Ferretti, F.; Pellegrino, G.; Raneri, M.; Schiavoni, M.; Caselli, A.; Briani, F. Sanguinarine Inhibits the 2-Ketogluconate Pathway of Glucose Utilization in Pseudomonas Aeruginosa. Front. Microbiol. 2021, 12, 2552.

(20) Sunthamala, N.; Suebsamran, C.; Khruaphet, N.; Sankla, N.; Janpirom, J.; Khankhum, S.; Thiwthong, R.; Chuncher, S. Sanguinarine and Chelidonine Synergistically Induce Endosomal Toll-like Receptor and M1-Associated Mediators Expression. J. Pure Appl. Microbiol. 2020, 14, 2351–2361.

(21) Mouton, J. M.; Heunis, T.; Dippenaar, A.; Gallant, J. L.; Kleynhans, L.; Sampson, S. L. Comprehensive Characterization of the Attenuated Double Auxotroph Mycobacterium TuberculosisΔleuDΔpanCD as an Alternative to H37Rv. Front. Microbiol. 2019, 10, 1922.

(22) Palomino, J.-C.; Martin, A.; Camacho, M.; Guerra, H.; Swings, J.; Portaels, F. Resazurin Microtiter Assay Plate: Simple and Inexpensive Method for Detection of Drug Resistance in Mycobacterium Tuberculosis. Antimicrob. Agents Chemother. 2002, 46 (8), 2720–2722.

(23) Ramirez, J.; Guarner, F.; Bustos Fernandez, L.; Maruy, A.; Sdepanian, V. L.; Cohen, H. Antibiotics as Major Disruptors of Gut Microbiota. Front. Cell. Infect. Microbiol. 2020, 10, 731.

(24) Willing, B. P.; Russell, S. L.; Finlay, B. B. Shifting the Balance: Antibiotic Effects on Host– Microbiota Mutualism. Nat. Rev. Microbiol. 2011, 9 (4), 233–243.

(25) Spaulding, C. N.; Klein, R. D.; Schreiber, H. L.; Janetka, J. W.; Hultgren, S. J. Precision Antimicrobial Therapeutics: The Path of Least Resistance? Npj Biofilms Microbiomes 2018, 4 (1), 1–7.

(26) Mathew, B.; Hobrath, J. V.; Ross, L.; Connelly, M. C.; Lofton, H.; Rajagopalan, M.; Guy, R. K.; Reynolds, R. C. Screening and Development of New Inhibitors of FtsZ from M. Tuberculosis. PloS One 2016, 11 (10), e0164100.

(27) Andreu, N.; Zelmer, A.; Fletcher, T.; Elkington, P. T.; Ward, T. H.; Ripoll, J.; Parish, T.; Bancroft, G. J.; Schaible, U.; Robertson, B. D.; Wiles, S. Optimisation of Bioluminescent Reporters for Use with Mycobacteria. PloS One 2010, 5 (5), e10777.

(28) Andreu, N.; Fletcher, T.; Krishnan, N.; Wiles, S.; Robertson, B. D. Rapid Measurement of Antituberculosis Drug Activity in Vitro and in Macrophages Using Bioluminescence. J. Antimicrob. Chemother. 2012, 67 (2), 404–414.

(29) Schaaf, K.; Hayley, V.; Speer, A.; Wolschendorf, F.; Niederweis, M.; Kutsch, O.; Sun, J. A Macrophage Infection Model to Predict Drug Efficacy Against Mycobacterium Tuberculosis. Assay Drug Dev. Technol. 2016, 14 (6), 345–354.

(30) Lee, N.; Nguyen, H. Ethambutol. In StatPearls; StatPearls Publishing: Treasure Island (FL), 2022.

(31) Paardt, A.-F. van der; Wilffert, B.; Akkerman, O. W.; Lange, W. C. M. de; Soolingen, D. van; Sinha, B.; Werf, T. S. van der; Kosterink, J. G. W.; Alffenaar, J.-W. C. Evaluation of Macrolides for Possible Use against Multidrug-Resistant Mycobacterium Tuberculosis. Eur. Respir. J. 2015, 46 (2), 444–455.

(32) Rastogi, N.; Goh, K. S.; Ruiz, P.; Casal, M. In Vitro Activity of Roxithromycin against the Mycobacterium Tuberculosis Complex. Antimicrob. Agents Chemother. 1995, 39 (5), 1162–1165.

(33) Falzari, K.; Zhu, Z.; Pan, D.; Liu, H.; Hongmanee, P.; Franzblau, S. G. In Vitro and in Vivo Activities of Macrolide Derivatives against Mycobacterium Tuberculosis. Antimicrob. Agents Chemother. 2005, 49 (4), 1447–1454.

(34) Dutta, N. K.; Karakousis, P. C. Latent Tuberculosis Infection: Myths, Models, and Molecular Mechanisms. Microbiol. Mol. Biol. Rev. MMBR 2014, 78 (3), 343–371.

(35) Grant, S. S.; Kawate, T.; Nag, P. P.; Silvis, M. R.; Gordon, K.; Stanley, S. A.; Kazyanskaya, E.; Nietupski, R.; Golas, A.; Fitzgerald, M.; Cho, S.; Franzblau, S. G.; Hung, D. T. Identification of Novel Inhibitors of Non-Replicating Mycobacterium Tuberculosis Using a Carbon Starvation Model. ACS Chem. Biol. 2013, 8 (10), 10.

(36) Gibson, S. E. R.; Harrison, J.; Cox, J. A. G. Modelling a Silent Epidemic: A Review of the In Vitro Models of Latent Tuberculosis. Pathog. Basel Switz. 2018, 7 (4), E88.

(37) Ravesloot-Chávez, M. M.; Van Dis, E.; Stanley, S. A. The Innate Immune Response to Mycobacterium Tuberculosis Infection. Annu. Rev. Immunol. 2021, 39, 611–637.

(38) Gopinath, V.; Raghunandanan, S.; Gomez, R. L.; Jose, L.; Surendran, A.; Ramachandran, R.; Pushparajan, A. R.; Mundayoor, S.; Jaleel, A.; Kumar, R. A. Profiling the Proteome of Mycobacterium Tuberculosis during Dormancy and Reactivation. Mol. Cell. Proteomics MCP 2015, 14 (8), 2160–2176.

(39) Pitts, B.; Meyerson, L. Inhibition of Na,K-ATPase Activity and Ouabain Binding by Sanguinarine. Drug Dev. Res. 1981, 1, 43–49.

(40) Bhusal, Y.; Shiohira, C. M.; Yamane, N. Determination of in Vitro Synergy When Three Antimicrobial Agents Are Combined against Mycobacterium Tuberculosis. Int. J. Antimicrob. Agents 2005, 26 (4), 292–297.

(41) Caleffi-Ferracioli, K. R.; Maltempe, F. G.; Siqueira, V. L. D.; Cardoso, R. F. Fast Detection of Drug Interaction in Mycobacterium Tuberculosis by a Checkerboard Resazurin Method. Tuberc. Edinb. Scotl. 2013, 93 (6), 660–663.

(42) Afriyie-Asante, A.; Dabla, A.; Dagenais, A.; Berton, S.; Smyth, R.; Sun, J. Mycobacterium Tuberculosis Exploits Focal Adhesion Kinase to Induce Necrotic Cell Death and Inhibit Reactive Oxygen Species Production. Front. Immunol. 2021, 12.

(43) Slaninová, I.; Táborská, E.; Bochoráková, H.; Slanina, J. Interaction of Benzo[c]Phenanthridine and Protoberberine Alkaloids with Animal and Yeast Cells. Cell Biol. Toxicol. 2001, 17 (1), 51–63.

(44) Sampson, S. L.; Dascher, C. C.; Sambandamurthy, V. K.; Russell, R. G.; Jacobs, W. R.; Bloom, B. R.; Hondalus, M. K. Protection Elicited by a Double Leucine and Pantothenate Auxotroph of Mycobacterium Tuberculosis in Guinea Pigs. Infect. Immun. 2004, 72 (5), 3031–3037.

