## Supplementary Figures S1-S4 for "Discovery of benzo[c]phenanthridine derivatives with potent activity against multidrug resistant *Mycobacterium tuberculosis*"

### Table of Contents for Supporting Information

Figure S1-S4

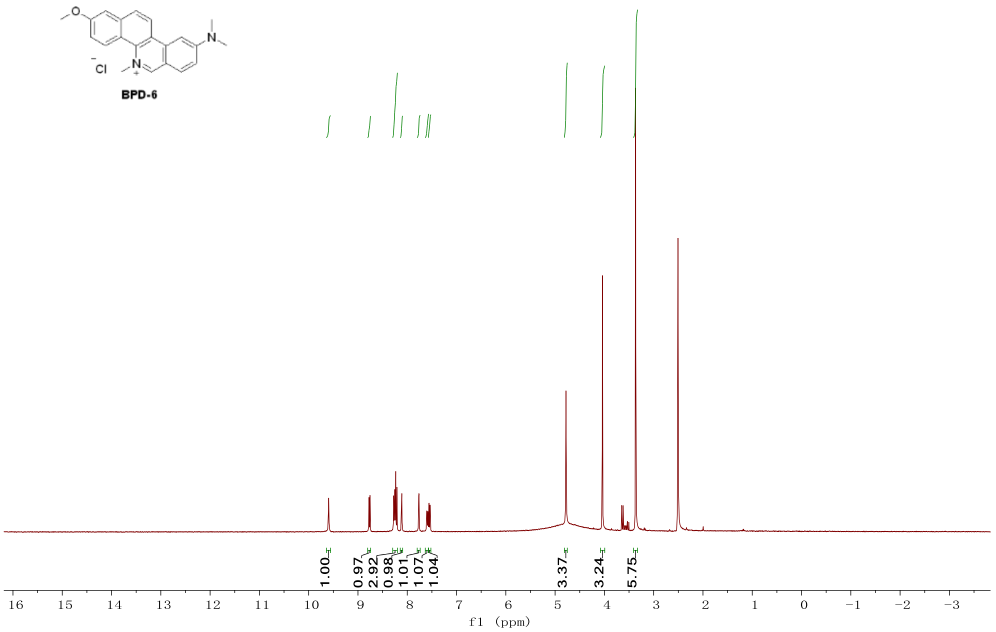


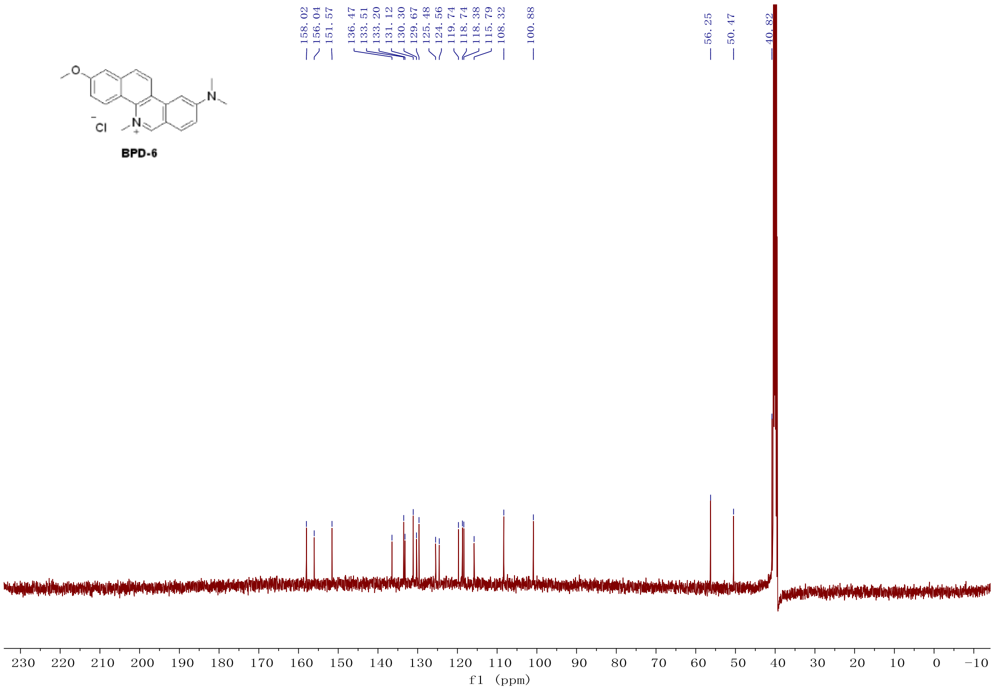


**Figure S1.** NMR spectra of 9-(dimethylamino)-2-methoxy-5-methylbenzo[c]phenanthridin-5-ium chloride (**BPD-6**): yellow solid, $^{1}$**H NMR** (400 MHz, DMSO-$d_6$): δ 9.59 (s, 1H), 8.77 (d, *J* = 9.0 Hz, 1H), 8.29 – 8.20 (m, 3H), 8.11 (d, *J* = 2.4 Hz, 1H), 7.77 (d, *J* = 2.3 Hz, 1H), 7.59 (dd, *J* = 9.3, 2.2 Hz, 1H), 7.55 (dd, *J* = 8.9, 2.3 Hz, 1H), 4.78 (s, 3H), 4.04 (s, 3H), 3.37 (s, 6H). $^{13}$**C NMR** (126 MHz, DMSO): δ 158.02, 156.04, 151.57, 136.47, 133.51, 133.20, 131.12, 130.30, 129.67, 125.48, 124.56, 119.74, 118.74, 118.38, 115.79, 108.32, 100.88, 56.25, 50.47, 40.82. **HRMS** (ESI) calcd. for $C_{21}H_{21}N_2O$ [M $^{+}$]: 317.1648, found: 317.1648. Yield for two steps 52%. Total yield 2.6%.

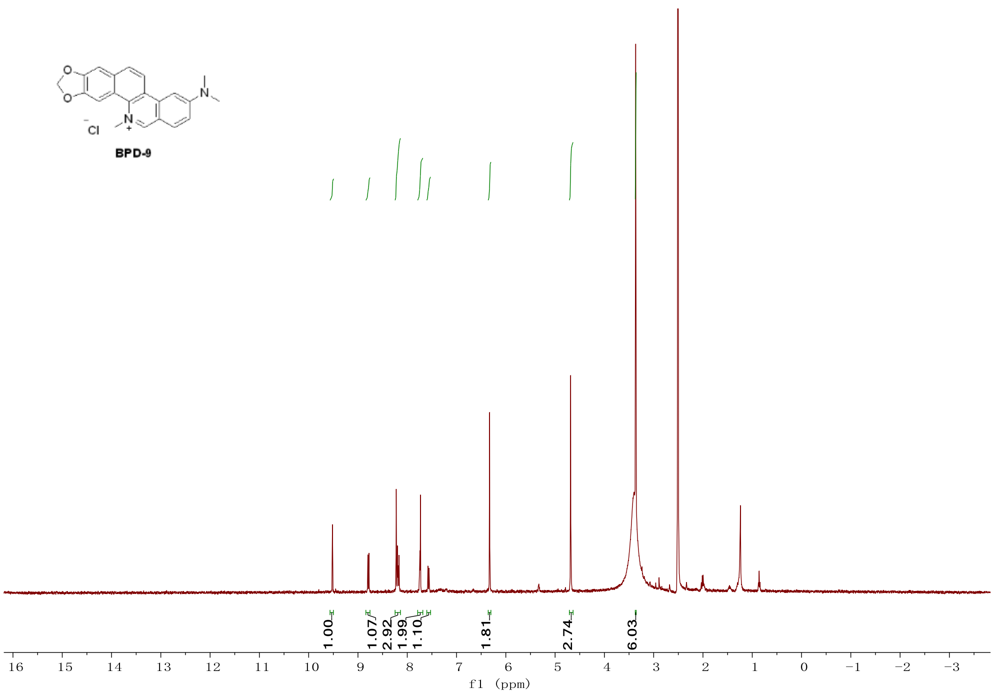


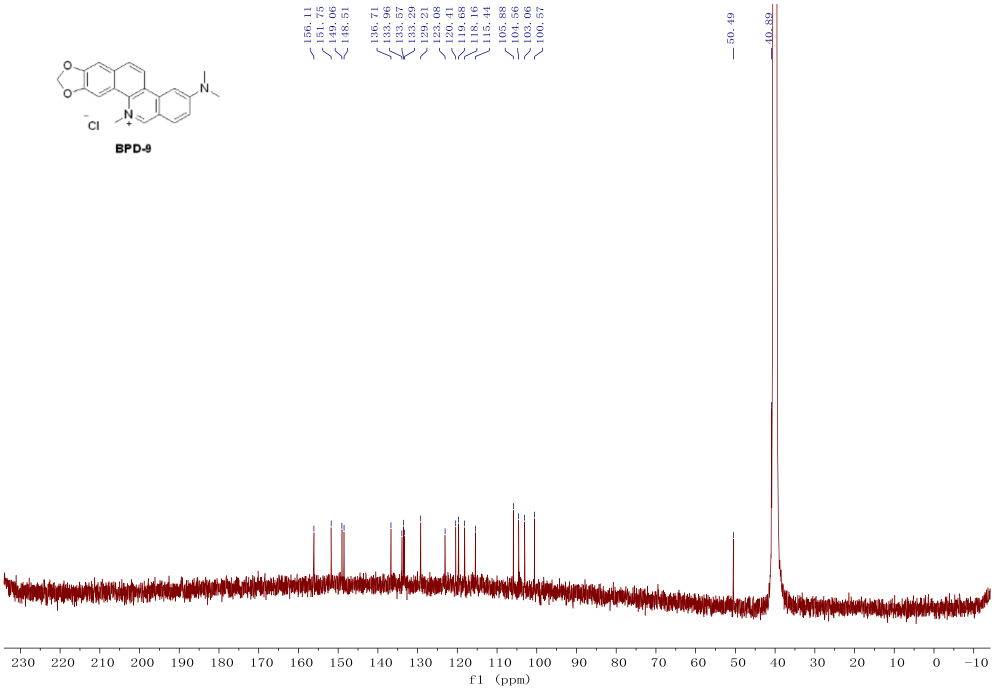


**Figure S2.** NMR spectra of 3-(dimethylamino)-12-methyl-[1,3]dioxolo[4',5':4,5]benzo[1,2-c]phenanthridin-12-ium chloride (**BPD-9**): yellow solid, $^{1}$**H NMR** (400 MHz, DMSO-$d_6$): δ 9.52 (s, 1H), 8.79 (d, $J$ = 9.0 Hz, 1H), 8.25 – 8.14 (m, 3H), 7.79 – 7.71 (m, 2H), 7.57 (dd, $J$ = 9.3, 2.2 Hz, 1H), 6.33 (s, 2H), 4.68 (s, 3H), 3.36 (s, 6H). $^{13}$**C NMR** (126 MHz, DMSO): δ 156.11, 151.75, 149.06, 148.51, 136.71, 133.96, 133.57, 133.29, 129.21, 123.08, 120.41, 119.68, 118.16, 115.44, 105.88, 104.56, 103.06, 100.57, 50.49, 40.89. **HRMS** (ESI) calcd. for $C_{21}H_{19}N_2O_2$ [M $^{+}$]: 331.1441, found: 331.144. Yield for two steps 46%. Total yield 1.2%.

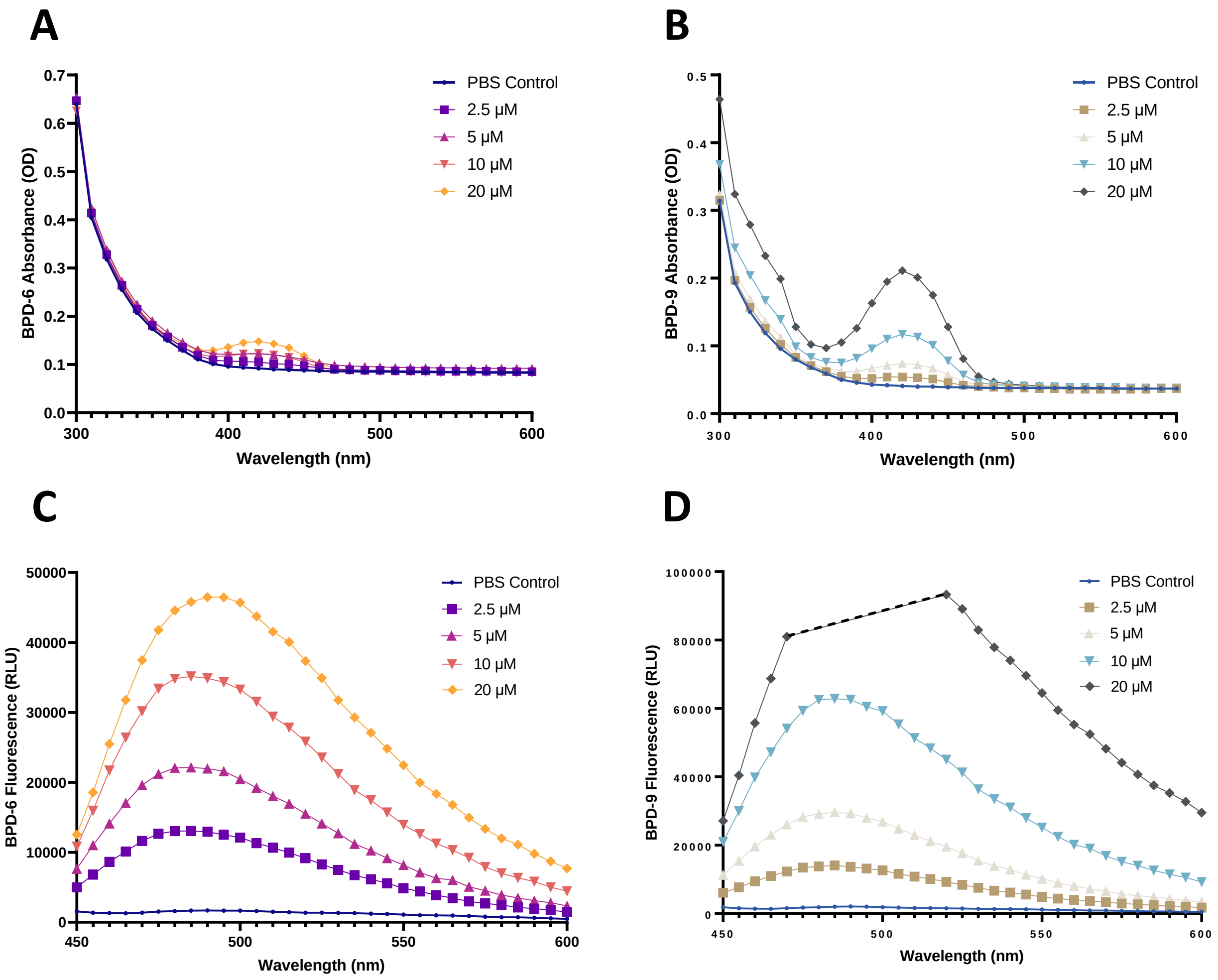


**Figure S3.** Absorbance and emission spectra scan of BPD-6 and BPD-9. Compounds were serially diluted (2.5 - 20 μM) in PBS buffer and the absorbance (A, B) or the emission (C, D) scan of BPD-6 and BPD-9 were measured. For the emission scan, the excitation wavelength was fixed at 420 nm.

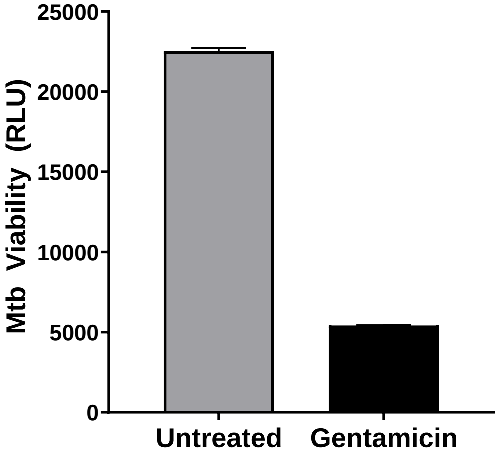


**Figure S4.** Relative viability of Mtb-*lux* as indicated by relative luminescence units (RLU). Mtb-*lux* was treated with 105 μM gentamicin or buffer alone for 24 h in PBS buffer.
